# A neural circuit for stereopsis in the fly

**DOI:** 10.64898/2026.11.11.765991

**Authors:** Duncan S. Mearns, Yoonwoo Park, Inwoo Shin, Mala Murthy

## Abstract

Stereopsis involves combining different views from the two eyes to infer depth, an ability that has intrigued neuroscientists for almost two centuries (Wheatstone, 1838). It enables insects such as the praying mantis to estimate the distance of prey (Nityananda et al., 2018), or primates to fixate objects at different depths (Cumming and Judge, 1986; Rashbass and Westheimer, 1961). While theoretical models for stereopsis have been proposed based on neural activity recordings (Marr and Poggio, 1976; Grossberg, 1983; Ohzawa et al., 1990), we do not have a detailed circuit mechanism for stereopsis founded on neural wiring. Here we achieve this in the fly, *Drosophila melanogaster*, by analyzing two whole-brain connectomes (Dorkenwald et al., 2024; Berg et al., 2026). Flies have sufficient overlap between the eyes to utilize binocular vision (Kemppainen et al., 2022); however, a circuit that performs stereopsis has not been uncovered in this model system. We show that a cell type that connects the two optic lobes called LC14 (Otsuna and Ito, 2006) forms a recurrent circuit across the brain that is capable of depth estimation. By building a connectome-constrained model, we demonstrate that recurrence is critical for binocular distance tuning in this circuit. Tuning is optimal for the size and distances of social targets (other flies), and we uncover a pathway within the brain specific for binocular vision computations. Silencing LC14 neurons renders this downstream circuitry sensitive to only monocular information and alters distance-dependent courtship behaviors, while activation of LC14 neurons causes flies to slow in response to an illusory object in their binocular visual field. In sum, we predict function from structure, uncovering how the fly brain computes distance from binocular cues, and identify a role for binocular vision in patterning social behavior.

## Results

Paired eyes allow animals to leverage the slightly different view of the world each sees to compute binocular disparities, the angular difference between an object’s location on the left and right eyes, which varies with distance. This computation involves coincidence detection; a circuit that performs stereopsis should integrate binocular information, amplify activity when objects fall at a preferred binocular disparity, and suppress activity when objects fall at all other non-preferred disparities via lateral inhibition (Marr and Poggio, 1976) (**Fig. 1A**). Neural activity recordings have uncovered binocular disparity-tuned neurons in cats (Barlow et al., 1967; Pettigrew et al., 1968), primates (Hubel and Wiesel, 1970), owls (Nieder and Wagner, 2000), mice (Scholl et al., 2013), tree shrews (Tanabe et al., 2022) and praying mantids (Rosner et al., 2019). Multiple possible circuit architectures could account for this tuning (**Fig. 1A**, bottom right). Prevailing models for both mammals and insects involve the feedforward integration of signals from monocular neurons with center-surround receptive fields onto binocular neurons in a higher visual area (Ohzawa et al., 1990; O’Keeffe et al., 2022); however, alternative models involving recurrent connectivity can also perform stereopsis (Grossberg, 1983), and have been speculated as a possible mechanism in insects (Rosner et al., 2019). Resolving which circuit architecture underlies the computation requires mapping the underlying neural connectivity (Briggman et al., 2011; Hubel and Wiesel, 1962; Reid, 2012), which is now possible in the fly, a premiere model for deciphering neural mechanisms of visual motion and feature detection (Borst and Groschner, 2023; Clark and Demb, 2016; Currier et al., 2023; Frye, 2015; Tanaka and Portugues, 2025). Fly connectomes contain neuronal morphologies (Dorkenwald et al., 2022), chemical synaptic connections (Buhmann et al., 2021; Yu et al., 2025), cell type annotations (Berg et al., 2026; Matsliah et al., 2024; Nern et al., 2025; Schlegel et al., 2024), and predicted neurotransmitter expression (Eckstein et al., 2024); however, interpreting how computation arises from this information still remains a challenge (Lin and Murthy, 2025; Litwin-Kumar and Turaga, 2019).

**Figure 1.**
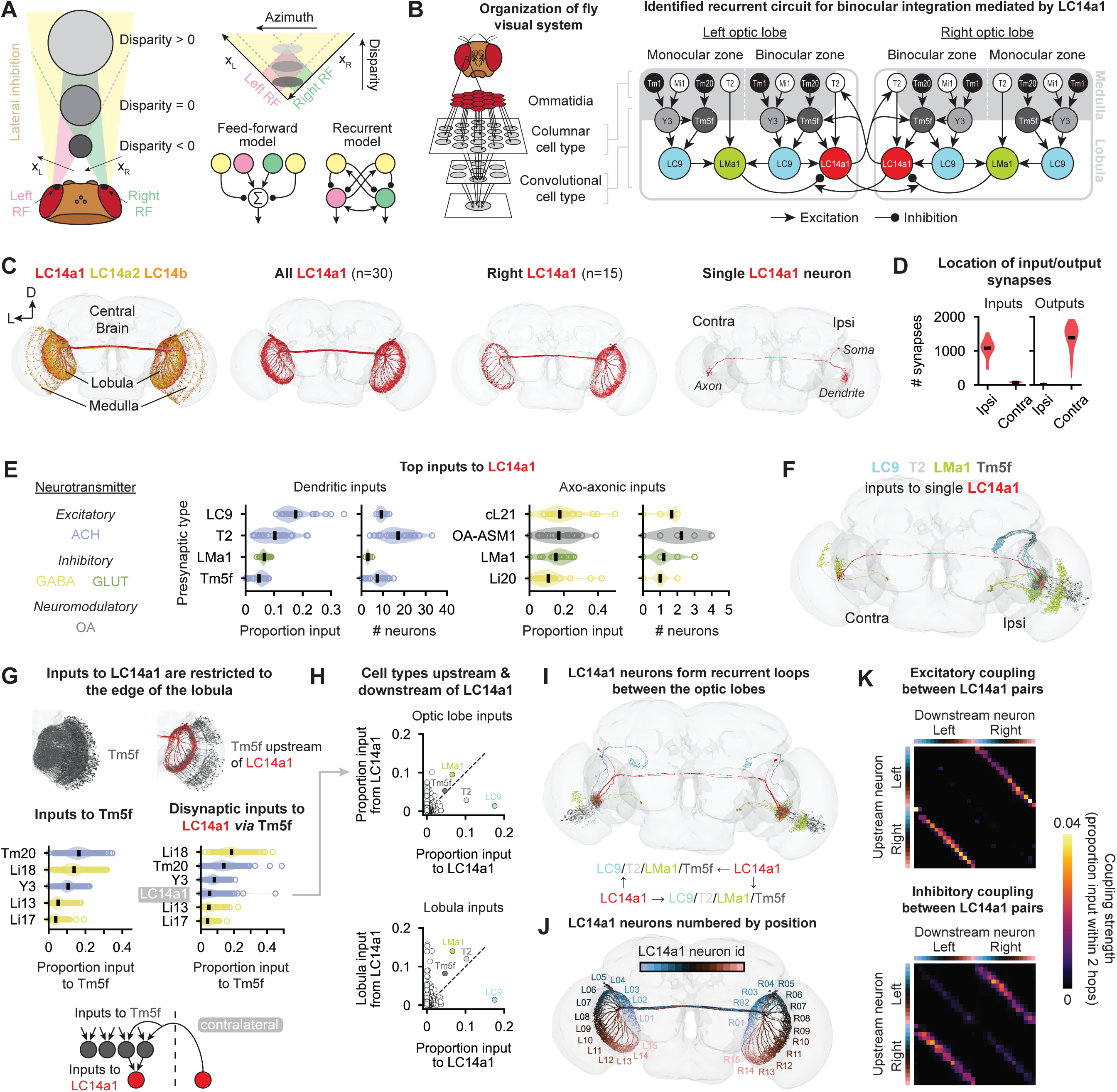
Recurrent circuitry for binocular integration via LC14a1 neurons. A. Distance estimation using stereopsis. Binocular disparity is defined as the difference in position (where the viewed object falls on the retina) between the two eyes relative to a reference point. Left: receptive fields (RFs) of neurons in each eye in the binocular visual field (magenta/green cones) intersect at a specific distance (disparity = 0). Lateral inhibition (yellow) suppresses responses to size-matched objects at other disparities. Stereopsis is used to discern a larger, further object from a smaller, closer one. Upper right: each pair of points in the left and right eye corresponds to a specific azimuth and disparity. Lower right: two models for computing distance from binocular distance cues. Feed-forward model: a binocular neuron sums excitatory and inhibitory inputs from each eye. Recurrent model: left and right neurons are reciprocally coupled. B. Schematic of a circuit for stereopsis identified in the fly optic lobes. Left: each eye consists of ∼800 ommatidia arranged in a hexagonal array. Ommatidia contain the photoreceptors and are in 1:1 correspondence with 31 columnar cell types in the medulla. Downstream cell types integrate information over columns, often combining information from multiple types with a convolutional architecture. Right: circuit for stereopsis identified in this study. Columnar types are shown at the top of the diagram, with subsequent types integrating information predominantly from previous layers. LC14a1 forms recurrent connections between the optic lobes in the binocular zone and receives lateral inhibition from the edge of the binocular zone and monocular zone via LMa1. C. Morphology of LC14 neurons. Left: all LC14 neurons from the FlyWire connectome, colored by subtype. Center left: all LC14a1 neurons (n=30). Center right: all LC14a1 neurons with their soma on the right (n=15). Right: morphology of a single LC14a1 neuron with soma on right. D. Number of input and output synapses on the ipsilateral and contralateral neurites of LC14a1 (n=30 neurons). Black bars = mean. E. Input structure of LC14a1 for strongest input types. Left: dendritic inputs. Right: axo- axonic inputs. Left subplots show proportion of input synapses coming from presynaptic cell types; right subplots show number of presynaptic partners of each type. Each point is a single LC14a1 neuron; black bars = mean. See **Fig. S1** for additional input types. F. A single LC14a1 neuron (red, soma on right) and all its LC9, T2, LMa1 and Tm5f presynaptic partners. G. Neurons upstream of LC14a1 receive feedback from contralateral LC14a1. Top left: all Tm5f neurons in the right optic lobe (gray); top right: only Tm5f neurons presynaptic to LC14a1 (red). Middle: inputs to Tm5f by cell type for all Tm5f neurons (left) vs. only Tm5f upstream of LC14a1 (right). Contralateral LC14a1 are top inputs to the subset of Tm5f presynaptic to LC14a1. Bottom: summary. H. LC14a1 are coupled via disynaptic connections through many cell types. Top: proportion of input neurons receive from LC14a1 vs. how much input they provide to LC14a1, grouped by cell type. Bottom: same analysis considering lobula inputs only. I. Two LC14a1 neurons (red) and their shared input and output partners (LC9: blue; T2: white; LMa1: green; Tm5f: gray) Bottom: information flow through circuit. J. LC14a1 tiles the edge of the lobula. Each LC14a1 on the left and right is given a unique index from one to 15 starting in the posterior visual field and sequentially increasing dorsally, anterior and then ventrally. K. Local recurrent coupling between LC14a1 neurons. Left and right LC14a1 neurons are ordered according to their topographic location in the lobula (see **J**). Coupling strength is the proportion of input within two hops upstream of a given neuron via excitatory or inhibitory intermediate neurons.

Flies are known to rely on a visual estimate of distance for a variety of behaviors, including evading predators (Card and Dickinson, 2008; Hammond and O’Shea, 2007), gap crossing (Pick and Strauss, 2005), and social interactions (Coen et al., 2016; Roemschied et al., 2023). While flies can use monocular cues – such as looming (de Vries and Clandinin, 2012), optic flow (Creamer et al., 2018) or motion parallax (Shomar et al., 2025) – to measure distance, binocular cues could augment flies’ distance estimation abilities. Although there is evidence that the *Drosophila* visual system has sufficient acuity to compute depth from binocular cues (Kemppainen et al., 2022), a circuit that performs binocular disparity detection has yet to be identified in the fly.

Each compound eye of the fly comprises a hexagonal array of 750-800 ommatidia spaced ∼5° apart (Zhao et al., 2025), which contain the photoreceptors. Photoreceptors that detect light from the same visual angle converge on downstream neurons in the lamina and medulla (Agi et al., 2014) (**Fig. 1B**, left). The medulla contains numerous columnar cell types that tile visual space and are in approximately one-to-one correspondence with specific locations in the visual field (Matsliah et al., 2024; Nern et al., 2025). Other cell types in the medulla and in downstream visual neuropils of the lobula and lobula plate integrate visual information from multiple medulla columns and tile the whole, or part, of the visual field. Thus the fly visual system resembles a convolutional neural network, with neurons in deeper layers integrating inputs from earlier layers. The structured organization of the *Drosophila* visual system has facilitated functional predictions from wiring (Borst, 2025; Hoeller et al., 2026; Lappalainen et al., 2024; Seung, 2024a; Ziyin et al., 2026), but so far these predictions have mostly ignored recurrent feedback or are based on large-scale simulations almost as detailed as the underlying circuitry. Here, we introduce a modeling framework that is simple enough to be interpretable, but complex enough to include all relevant cell types and their polysynaptic and recurrent connections from the connectome. Importantly, we do not know the function of this network *a priori*; we learn it directly from the wiring. Our model of the recurrent circuitry between the eyes in the binocular visual field makes specific, testable predictions, and, by leveraging knowledge of downstream cell types, we link this circuitry to ethologically relevant behaviors.

### A recurrent circuit combines information from the two eyes

LC14 neurons in *Drosophila* (Hassan et al., 2000; Linneweber et al., 2020; Otsuna and Ito, 2006) have been proposed as a potential substrate for stereopsis (Fenk et al., 2022; Kemppainen et al., 2022; Rosner et al., 2019) due to their morphological similarity to binocular disparity tuned neurons in the praying mantis, the only insect to date known to perform stereopsis (Read, 2021). There are three subtypes of LC14 in the connectome that differ in connectivity: LC14a1, LC14a2 and LC14b (Matsliah et al., 2024; Nern et al., 2025) (**Fig. 1C**). Each subtype connects the two optic lobes, with neurites restricted to the edge of the lobula, corresponding to the edge of the visual field seen by each eye (**Fig. 1C**). The majority of input synapses (94.0 ± 2.1%, mean ± std) occur on the neurite ipsilateral to the cell body (which we call the dendrite) and outputs are in the contralateral neurite (which we call the axon) (**Fig. 1D**). We focus our analysis on LC14a1 for two reasons: i) its connections support a circuit for stereopsis as we demonstrate below and ii) this subtype is connected to downstream circuitry relevant for male courtship behavior, which we use to interpret circuit function (see **Fig. 4**). LC14b participates in stereovision but has a different function from LC14a1 (see **Fig. 4**), while LC14a2 may play a role in detecting movements of the eyes (see **Discussion**).

We use the number of synaptic connections between neurons as a proxy for synaptic strength (Liu et al., 2022) (see **Methods**). Four cell types account for 40% of the dendritic input to LC14a1 (**Fig. 1E** & **Fig. S1**). Three of these cell types (LC9, T2 and Tm5f) are cholinergic (excitatory) and one, LMa1, is glutamatergic. Glutamate can act as an inhibitory neurotransmitter in insects (Liu and Wilson, 2013; Mauss et al., 2015) and is expected to be inhibitory in this circuit due to the expression of glutamatergic-gated chloride channels in LC14 neurons (*Y. Kermangaliev, personal communication*). Both LC9 and T2 are small object detectors (Aptekar et al., 2015; Keleş et al., 2020), and also represent the top inputs to LMa1 (**Fig. S1**). Such a motif suggests that LMa1 provides normalizing inhibition to LC14a1 (Carandini and Heeger, 2012; Seung, 2024b). LMa1 also makes a small number of axo-axonic synapses onto LC14a1 (**Fig. 1E**, right). Morphologically, the LMa1 inputs to a given LC14a1 neuron cover a larger area of the lobula than the excitatory inputs (**Fig. 1F**), providing a mechanistic basis for lateral inhibition necessary for stereopsis (Marr and Poggio, 1976), which we quantify below (see **Figs. 2** & **3**).

**Figure 2.**
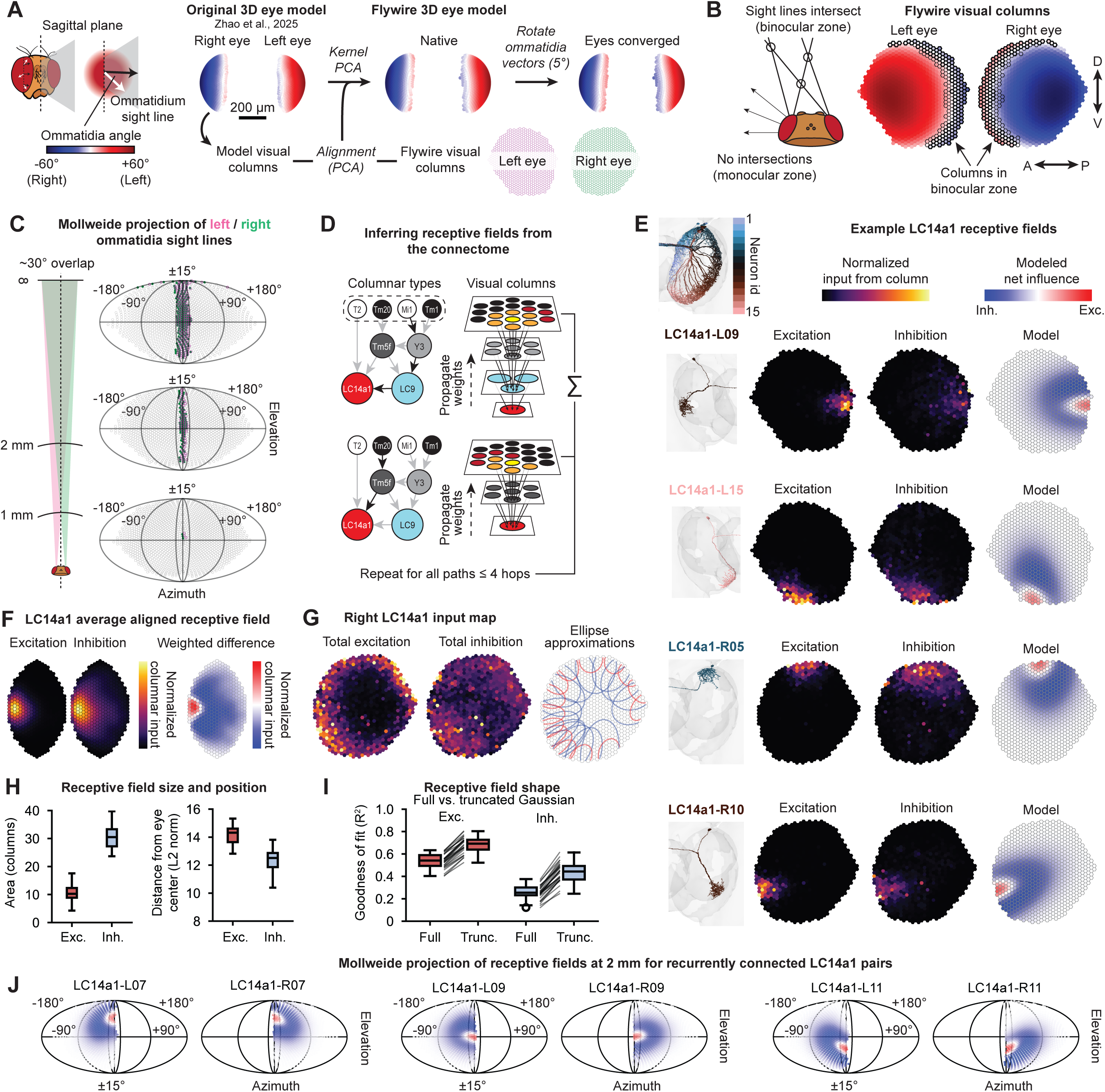
Excitatory and inhibitory receptive fields of LC14a1 neurons. A. Left: each ommatidium has a 3D position on the surface of the fly eye and points in a direction that varies smoothly over the eye. We define the ommatidia angle as the angle between the viewing direction of an ommatidium and the sagittal plane. Positive angles indicate an ommatidium is viewing the left visual hemifield and negative angles indicate an ommatidium is viewing the right visual hemifield. Center left: Location and angle of ommatidia from Zhao et al. (2025). Center right: virtual eye model for the FAFB-FlyWire fly after mapping visual columns from the connectome to theoretical positions and viewing angles using kernel PCA. Right: viewing angles of the virtual FAFB-FlyWire eye after converging ommatidia 5°. B. Neural columns that can contribute to stereovision. When the eyes are maximally converged, sightlines of ommatidia from one eye intersect with the sightlines of the other eye at different distances. Any ommatidium/column whose sightline intersects with those of the contralateral eye could be used to compute disparity of objects and are outlined in black. Dark outline = binocular columns when eyes are in native position (127/785, left, 133/796, right); light outline = columns that become binocular when eyes converge (192/785, left, 194/796, right). Neural columns are shown as if looking through the fly’s eyes (i.e., reflected relative to A). C. Projection of ommatidia sightlines form the left and right eye at different distances. Below 1 mm sight lines from the left and right eye do not intersect. Overlap is maximal at infinity and covers ∼30° of the visual field. Ommatidia that intersect with sightlines from the contralateral eye at each distance are colored (magenta = left eye, green = right eye). D. Mapping receptive fields of LC14a1 neurons. Input weights (proportion of synapses) are propagated back through the connectome to columnar types (see Fig. 1B) via all excitatory neurons within four hops. Input maps resulting from each path are summed to compute the final contribution of each visual column to the receptive field of a given LC14a1 neuron. For inhibitory receptive fields, excitatory paths to inhibitory neurons one hop upstream of LC14a1 are computed (see **Methods**). E. Inferred receptive fields of four example LC14a1 neurons. Each row shows the neuron morphology (left), normalized excitatory input from columns (center left), normalized inhibitory input from columns (center right), and modeled difference of gaussian receptive field (right). See **Fig. S3** for all neurons. F. Average excitatory and inhibitory receptive fields for LC14a1 neurons. Left: average excitatory and inhibitory inputs from columns after aligning all individual receptive fields to the edge of the visual column map. Right: difference between excitatory and inhibitory columnar input relative to edge of visual column map. G. Activation maps for all LC14a1 in the right optic lobe. Left: total excitatory columnar input. Center: total inhibitory columnar input. Right: overlaid ellipse approximations for all individual excitatory and inhibitory receptive fields (red = excitatory, blue = inhibitory). H. Size (left) and position (right) of excitatory and inhibitory receptive fields. Size is area of a fitted ellipse; distance is Euclidean distance of the receptive field centroid from the center of the eye. I. Model fit for full and truncated 2D Gaussians for the excitatory and inhibitory receptive fields. Lines show difference in goodness of fit (R^2^) between modeled columnar input strength for each neuron. J. Receptive fields projected through ommatidia sight lines to a distance of 2 mm for three reciprocally coupled pairs of LC14a1 neurons.

**Figure 3.**
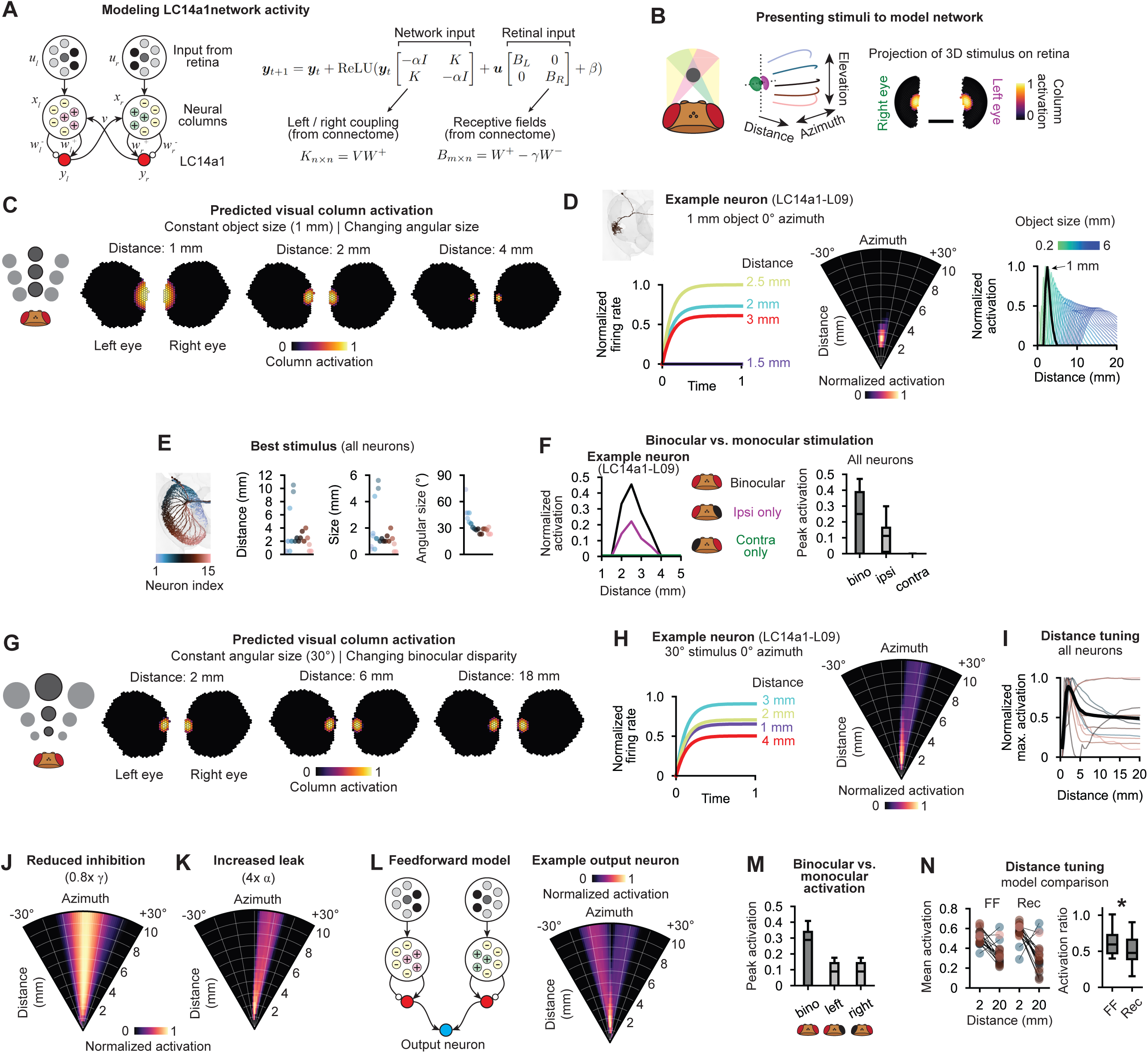
A connectome-constrained model of binocular integration. A. Connectome-constrained model of stereovision in *Drosophila*. Modeled as a two layer network with inputs coming from the retina (*u*) to visual columns (*x*). LC14a1 neurons (*y*) are modeled as leaky integrators with the strength of excitatory and inhibitory inputs from visual columns and the organization of recurrent inputs from other LC14a1 neurons are obtained from the connectome (see Figs 1 & 2). B. Presenting visual stimuli to the model. Spherical objects are placed at different azimuths, elevations and distances from the model fly and projected onto the retina to compute the input to the network. Scale bar = 200μm. C. Investigating size tuning in LC14a1 neurons. Projection of a 1 mm sized object at different distances onto the retina. Left: schematic showing objects of a constant angular size at different distances (not to scale). Right: column activation to a 1 mm object at different distances. D. Example LC14a1 response. Left: activity over time for a single LC14a1 neuron (L09) to a 1 mm-sized object in the center of the visual field at different distances. Center: maximum response to a 1 mm object placed at different azimuths and distances. Right: distance tuning to objects of different sizes. Each line is a tuning curve for an object of a different size. Black line = peak response tuning curve (1 mm object). See **Fig. S5** for all neurons. E. Stimulus that elicited the strongest response for each modeled neuron. Far left: neurons ordered and colored by index. Plots show the distance (left), size (center) and angular size (right) of the stimulus that elicited the strongest response. F. Non-linear amplification of binocular inputs. Left: response to a 1 mm object at different distances in the center of the visual field for a single LC14a1 neuron (L09) when inputs are provided to both or only one eye. Right: peak activation to a 1 mm object when the stimulus is viewed by both or one eye. Ipsi = ipsilateral, contra = contralateral eye with respect to LC14a1 neuron. G. Investigating binocular disparity tuning in LC14a1 neurons. Projection of a 30° object at different distances onto the retina. Left: schematic showing that as an object moves further away it must get larger to maintain constant angular size (not to scale). Right: column activation to a 30° object at different distances. H. As in (D), showing the same LC14a1 neuron response to an object of constant angular size at different distances. See **Fig. S6** for all neurons. I. Binocular distance tuning in LC14a1 neurons. Each colored line shows the maximum activation of a neuron to a 30° object at each distance, normalized to the maximum. Black line = population mean. J. Distance estimation depends on tuned inhibition. Same neuron as panels D & H. Activity elicited when the model parameter controlling inhibition strength () is reduced by 20%. K. Distance estimation depends on tuned leak current. Same neuron as panel D & H. Activity elicited when the model parameter controlling leak current is increased fourfold. L. Comparison to a feedforward model for stereopsis. Recurrent feedback is removed and the outputs of left and right LC14a1 neurons are integrated in a downstream neuron. Left: model schematic; right: example neuron activation to a 30° object. M. As (F), showing feedforward model responses to a 30° stimulus located at the preferred distance of each neuron when inputs are provided to one or both eyes. N. Comparison of distance tuning between the feedforward (FF) and recurrent (Rec) model. Left: mean activation of each neuron to a 30° stimulus placed at 2 mm and 20 mm. Lines connect responses of the same neuron. Right: decrease in responses with distance as measured by activation ratio (20 mm / 2 mm) for each model. * p < 0.05 Wilcoxon signed-rank test.

While all the primary input types to LC14a1 tile the visual field, only a subset of neurons of each type provides input to LC14a1 due to the restricted location of LC14a1 dendrites at the edge of the lobula (**Fig. 1G**, top). We find that the contralateral LC14a1 neurons are a major input to the neurons upstream of LC14a1 (**Fig. 1G,H**). Left and right LC14a1 neurons are disynaptically coupled via multiple interneuron types, creating a recurrent circuit between the optic lobes (see **Fig. 1B**). These recurrent connections between pairs of LC14a1 neurons are spatially restricted in the lobula and only occur between neurons viewing similar elevations of the visual field (**Fig. 1I- K**), indicating that the circuit performs local binocular computations, rather than broadly integrating signals across visual hemifields. Excitatory recurrent connections are stronger and more spatially restricted than inhibitory recurrent connections (**Fig. 1K**), providing a mechanism for preferential enhancement of objects falling within the excitatory receptive field of pairs of LC14a1 neurons, and suppression by objects misaligned in elevation.

This circuit has the hallmarks of a circuit for stereopsis: local recurrent excitation between locations on the two eyes corresponding to a given binocular disparity, with inhibition from locations corresponding to other binocular disparities. The recurrent architecture differs from the structure assumed to underlie early binocular disparity processing in mammals, which begins with the feed-forward integration of monocular signals from each eye (Cumming and DeAngelis, 2001; Ohzawa et al., 1990), but resembles alternative theoretical models for stereopsis that were proposed over 50 years ago (Grossberg, 1983, 1976).

### LC14a1 neurons have center-surround receptive fields in the binocular visual field

To compute disparity, reciprocally coupled LC14a1 neurons should have aligned receptive fields in the binocular zone, with focal excitatory centers and an inhibitory surround (**Fig. 1A**). The excitatory center would allow for coincidence detection of objects landing at a fixed binocular disparity, and the inhibitory surround would suppress responses to objects at other disparities. To test this, we leveraged a recently published 3D microCT scan of a fly head which contains the location and viewing direction of all the ommatidia of a female fly (Zhao et al., 2025). We aligned the visual columns from each brain connectome (Berg et al., 2026; Dorkenwald et al., 2024) to those of the microCT fly to infer the locations and viewing directions of the sensors that provide input to visual columns in the connectome (**Fig. 2A**; see **Methods**). Our initial mapping generated a small amount of binocular overlap (∼15°), less than the 20-30° known from experimental measurements (Kemppainen et al., 2022). This discrepancy can be accounted for by the fact that flies can move their photoreceptor array underneath their lenses, changing the viewing angle of their ommatidia (Falt et al., 2026; Fenk et al., 2022), so we modestly converged the ommatidia sightlines by 5° to bring the size of the binocular zone in line with experimental measurements (**Fig. 2A**, right).

Next, we identified which visual columns in the connectome view the binocular zone (**Fig. 2B**); the greatest binocular overlap occurred at anterior elevations of the visual field with 7-8 columns per eye contributing to binocular vision, and this overlap dropped to about 3 columns per eye in the dorsal and ventral parts of the visual field. We projected the viewing direction of each visual column into the world and found that different columns from each eye align at different distances, which correspond to different binocular disparities (**Fig. 2C**).

To infer the receptive fields of LC14a1 neurons, we mapped inputs back to visual columns (**Fig. 2D**, see **Methods**). This approach estimates the influence of a given visual column on an LC14a1 neuron. We estimated the excitatory (“classical”) and inhibitory (“extra-classical”) receptive fields separately, which revealed a consistent pattern of small, focal excitation from the binocular zone onto each LC14a1 neuron and a broader inhibitory surround extending into the monocular zone (**Fig. 2E-H** & **Fig. S3**). We found that LC14a1 receptive fields were best modeled as a difference of half-gaussians (**Fig. 2I**), with a focal excitatory receptive field situated more towards the edge of the visual field than the broader inhibitory surround.

Subtracting the inhibitory receptive field from the excitatory receptive field generates a final estimate of the net influence of a given column onto an LC14a1 neuron (**Fig. 2E**, right). Finally, projecting receptive fields through the sightlines of ommatidia in our 3D eye model revealed that reciprocal pairs of LC14a1 neurons view the same point in space, centered around the midline, ∼2 mm in front of the fly (**Fig. 2J**).

Thus, the receptive fields of LC14a1 lie predominantly within the binocular zone with excitation weighted towards the edge of each eye’s visual hemifield and recurrently connected pairs viewing the same region of visual space. These properties confer the network with all the requirements to generate binocular disparity tuning.

### A connectome-constrained computational model of stereopsis in the fly visual system

To investigate distance tuning in LC14a1 neurons, we built a connectome-constrained rate-based dynamics model of the network, with input from the 3D model of the fly eye (**Figs. 3A-B** & **Fig. S4,** see **Methods**). We modeled LC14a1 neurons as leaky integrators that receive weighted input from visual columns according to their excitatory and inhibitory receptive fields inferred from the connectome (see **Fig. 2**). Additionally, LC14a1 receives recurrent feedback from other LC14a1 neurons in the network as measured in the connectome (see **Fig. 1**).

We first determined to which angular size LC14a1 neurons are tuned (**Fig. 3C-F**), then subsequently held objects at a constant angular size on the retina to investigate binocular disparity tuning (**Fig. 3G-J**). As an object moves further away from the fly, its angular size appears smaller on the retina (**Fig. 3C**). Each stimulus we present drives an LC14a1 neuron to a different steady- state firing rate (**Fig. 3D**, left; one example LC14a1 neuron (L09) shown here). This allows us to generate a tuning curve for azimuth and distance for each object diameter (**Fig. 3D**, center & **Fig. S5**), revealing strong angular size tuning. Over all object diameters and distances tested, we found that almost all LC14a1 neurons (except for those in the posterior visual field) consistently preferred 1 mm diameter objects placed 2 mm away and centered in the visual field, corresponding to an angular size of ∼30° (**Fig. 3E** & **Fig. S4**). As expected from their morphology, LC14a1 neuron tuning curves tiled elevations in the center of the visual field, with finer resolution in the anteroventral region (**Fig. S4**). The anteroventral visual field corresponds to a functional “love- spot” in the male fly visual system, which contains a higher proportion of sexually dimorphic circuitry (Berg et al., 2026). LC14a1 neurons, however, have similar morphologies in the male and female connectomes (**Fig. S1**), although these neurons are predicted to express the sex determination transcription factor *fruitless* ((Deutsch et al., 2025) and *Y. Kurmangliev personal communication*), indicating sex-specific functions (see **Fig. 5** & **Discussion**).

To encode binocular depth information, neurons underlying stereovision should exhibit non-linear summation of the inputs from each eye (Marr and Poggio, 1976; Read, 2022). We tested responses to stimuli while blocking inputs to one eye in the model. Model neurons showed no response when only the contralateral eye could see the stimulus, but responded with roughly twice the amplitude to a binocular stimulus versus a stimulus seen in only the ipsilateral eye (**Fig. 3F**). This suggests that, in our model, inputs to the ipsilateral eye effectively “gate” the response to a stimulus in the contralateral eye.

Next, we held angular size constant and presented objects at different distances, ensuring that binocular disparity is the only cue available to the model (**Fig. 3G**). Even when angular size was held constant, LC14a1 neurons could discriminate objects at different depths (**Fig. 3H-I**, **Fig. S6**). We found that the model’s ability to compute depth from binocular cues depended largely on only two parameters: either reducing inhibition or increasing the leak current in the model impaired distance tuning (**Fig. 3J,K**). Our estimation of inhibition is reasonable because inhibitory synapses are typically stronger than excitatory synapses in fly visual circuits (Ammer et al., 2023). The leak parameter in our model controls the balance of LC14a1 neurons’ intrinsic time constant and the strength of recurrent feedback. A small leak allows for better integration of inputs, while a large leak reduces the ability of the circuit to perform integration (similar to models of the *Drosophila* central complex or oculomotor integrator of fish (Duan et al., 2025; Hulse and Jayaraman, 2020; Khona and Fiete, 2022; Kim et al., 2017; Lyu et al., 2022; Seung, 1996; Turner-Evans et al., 2017; Westeinde et al., 2024)).

Since most models of stereopsis are feed-forward (Ohzawa et al., 1990; O’Keeffe et al., 2022), we investigated how the recurrently connected model performs in comparison (see **Methods**). The feed-forward model addresses how a hypothetical neuron that sums the outputs of pairs of LC14a1 would compute depth compared to the recurrent architecture identified in the connectome (cf. **Fig 3A, L**). Such a feed-forward model does not produce the same non-linear amplification to binocular stimuli observed in the recurrent model (**Fig. 3M**). The recurrent model is also better at encoding depth, showing both stronger responses to near stimuli and weaker responses to far stimuli (**Fig. 3N**).

Our model, constrained primarily with weights measured from the connectome, and presented with stimuli as the fly would view them, predicts that LC14a1 neurons perform non-linear integration of inputs to the left and right eyes. Distance tuning in an LC14a1 neuron arises when it pushes its contralateral counterpart over threshold, initiating a recurrent feedback loop that amplifies responses only when an object falls within the excitatory receptive fields of both neurons. Lateral inhibition restricts the range of locations where this amplification is triggered. Given the interocular distance in our model of 0.68 mm, and preferred distance of ∼2 mm, we predict LC14a1 is tuned to a headcentric binocular disparity of ∼20°, i.e. an offset of 10° from the sagittal plane in each eye; however, the exact tuning would depend on the strength of lateral inhibition and the convergence angle of the eyes (**Fig S4**, see **Discussion**). Our model finds that all LC14a1 neurons in the anterior visual field are tuned to the same binocular disparity, in contrast to mammalian brains which contain a disparity map encoding multiple depths. Our results further suggest that detecting objects at 2 mm - about a body length away - is ethologically relevant (see below).

### Binocular distance information is relayed to a dedicated downstream pathway capable of stereopsis over a range of azimuths

If flies utilize binocular vision for distance estimation, information from LC14a1 should be routed to particular downstream cell types to drive distance-dependent behaviors. We searched among visual projection neurons (VPNs), the bottleneck between the optic lobes and the central brain (Cowley et al., 2024; Klapoetke et al., 2022) and which include other lobula columnar (LC) types, for those that receive LC14a1 input (**Fig. 4A**). Three VPN types emerged from this analysis: LC43, LC31 and LC10a; which are all predicted to be sexually dimorphic or connect strongly to sexually dimorphic circuitry (Berg et al., 2026; Deutsch et al., 2025). Of these, LC10a has the most similar input structure to LC14a1, indicating it responds to similar visual features. LC10a is known to detect conspecifics and to drive social behaviors such as courtship and aggression (Hindmarsh Sten et al., 2021; Ribeiro et al., 2018; Schretter et al., 2025, 2021), but its role in binocular computations has not been investigated.

**Figure 4.**
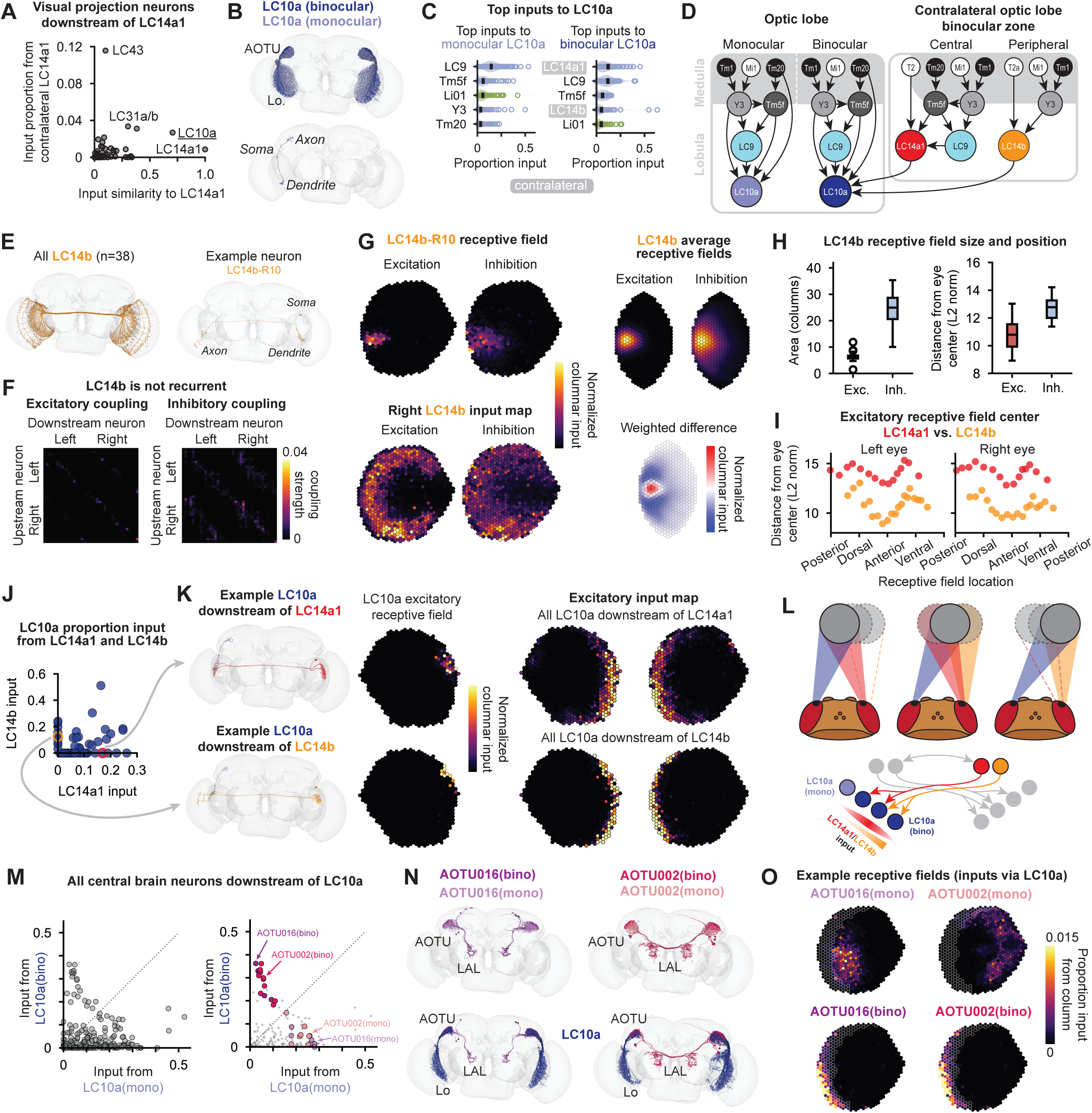
Downstream pathways for processing binocular information over a range of azimuths. A. Visual projection neurons vary in both their input similarity with and input from LC14a1 neurons. Input from contralateral LC14a1 plotted against input (cosine) similarity to LC14a1 for each visual projection neuron type. Each point is the mean per cell type. B. Morphology of LC10a neurons. Top: all LC10a neurons. Bottom: single LC10a neuron morphology. Monocular (light blue) = no input from contralateral eye via LC14; binocular (dark blue) = receives input from contralateral eye via LC14. C. Monocular and binocular LC10a neurons differ in their top input cell types. Left: top inputs to monocular LC10a. Right: top inputs to binocular LC10a neurons. Each point is an individual LC10a neuron. Black bar = mean. Blue = cholinergic input, green = glutamatergic input. D. Schematic of binocular integration in LC10a. LC10a in the binocular zone receive input from the contralateral eye via LC14a1 and LC14b. E. LC14b morphology. Left: all LC14b neurons. Right: single LC14b neuron (soma on the right). F. LC14b neurons are not recurrently coupled across the midline. Proportion input each LC14b neuron receives for every other LC14b neuron within two hops via excitatory (left) and inhibitory (right) paths. Cf. Fig. 1K. G. LC14b receptive fields. Top left: Single LC14b receptive field inferred from the connectome showing distribution of excitation (left) and inhibition (right) from visual columns. Bottom left: excitatory and inhibitory input map of all LC14b neurons with soma on the right. Top right: average excitatory and inhibitory columnar input to LC14b neurons aligned to the edge of the visual field. Bottom right: difference between average excitatory and inhibitory receptive fields. Cf. Fig. 2E**-G**. H. Size (left) and position (right) of LC14b excitatory and inhibitory receptive fields. Size is area of a fitted ellipse; distance is Euclidean distance of the receptive field centroid from the center of the eye. I. Excitatory receptive field center for all LC14a1 (red) and LC14b (orange) neurons. Each point is a neuron, ordered by anatomical location, showing the distance of the receptive field from the center of the eye. J. Proportion of input to individual binocular LC10a neurons from LC14a1 and LC14b. Each point is a single LC10a neuron. K. Receptive fields of LC10a neurons downstream of LC14a1 (top row) and downstream of LC14b (bottom row). Left: example LC10a neurons, with soma on the left, and their upstream LC14a1/LC14b partners (soma on the right). Center: excitatory receptive field of each example neuron. Right: excitatory input map for all LC10a neurons downstream of LC14a1 or LC14b. L. Schematic of binocular integration in binocular LC10a. Blue shading: receptive fields of three LC10a neurons; red shading: LC14a1 receptive field; orange shading: LC14b receptive field. Grey circles indicate objects at the same distance but different azimuthal locations detected by different LC10a. M. Integration of inputs from monocular and binocular LC10a downstream in the central brain. Each point is a single neuron in the central brain. Monocular and binocular LC10a input is the proportion of each neuron’s total input that comes from the given LC10a population. Dotted line indicates equal input from monocular and binocular LC10a. N. Central brain cell types receiving preferential input from binocular LC10a. Top row: morphology of all neurons of the given type. Bottom row: binocular neurons of the given type and all their presynaptic LC10a partners. Left (purple): AOTU016; right (pink): AOTU002. AOTU: anterior optic tubercle; LAL: lateral accessory lobe. O. Receptive fields of monocular and binocular central brain neurons. Weighted sum of LC10a excitatory receptive fields that provide input to a monocular neuron (top) and binocular neuron (bottom) for AOTU016 (left) and AOTU002 (right). White outlined columns indicate the binocular zone.

LC10a dendrites tile the entirety of the lobula (**Fig. 4B**); however, since LC14a1 synapses are only present at the edge of the lobula, only a subset of LC10a can receive binocular input. Although LC14a1 provides ∼2% of the input to LC10a as a population, when considering only the LC10a that receive LC14a1 input, LC14a1 becomes the strongest input, sometimes accounting for up to 40% of the synapses onto an LC10a dendrite (**Fig 4C**, right).

Parsing LC10a into binocular and monocular subsets reveals that binocular LC10a neurons also receive input from LC14b (**Fig. 4C,D**), which, similar to LC14a1, tiles the edge of the lobula (**Fig 4. D-E**). Unlike LC14a1, however, LC14b neurons do not participate in recurrent loops between the optic lobes (**Fig. 4F**), so on its own, LC14b likely does not compute depth (**Fig. 3**). However, LC14b has a center-surround receptive field, which is slightly inset from the edge of the visual column map compared to LC14a1 (**Fig. 4G,H** & **Fig. S7).** That is, LC14b receptive fields are shifted about 3 columns towards the center of the eye relative to LC14a1 (**Fig. 4I**). This corresponds to the edge of the binocular visual field on the ipsilateral side, i.e. the location at which an object first appears in the visual field of the *contralateral* eye (see **Fig 2B**).

LC14b thus appears to enable the binocular circuitry to perform stereopsis at a fixed disparity over a range of azimuths: solving the problem of maintaining responses in the binocular circuitry when an object slips to the left or right of the center of the binocular zone. A binocular LC10a neuron (70/234) can receive input solely from LC14a1 (43/70), solely from LC14b (9/70), or from both (18/70) (**Fig. 4J**). We computed the receptive fields of these different subsets of LC10a and found that they are topographically organized (**Fig. 4K** & **Fig. S7**): LC10a downstream of LC14b have their receptive fields at the innermost edge of the visual hemifield (i.e., viewing the greatest extent into the contralateral hemifield) where it would intersect with the receptive field of the LC14b in the contralateral eye (**Fig. 4L**, right); LC10a downstream of LC14a1 have their receptive fields in the center of the binocular zone, matching an LC14a1 in the contralateral eye (**Fig. 4L**, left). LC10a that receive input from both LC14b and LC14a1 have receptive fields intermediate to the other two subsets (**Fig. 4L**, middle). In short, LC10a neurons with receptive fields at different azimuthal locations receive input from contralateral LC14 with a complementary angular offset, and thus encode a fixed binocular disparity over a range of azimuths.

While most downstream targets of LC10a receive input predominantly from monocular LC10a (**Fig. 4M-N**), in both the male and female connectomes, we identified a number of neurons that receive input predominantly from binocular LC10a; these neurons belong primarily to only two cell types: AOTU016 and AOTU002 (**Fig. 4M**, right & **Fig. S7**). This organization was unique to LC10a and was not present in other LCs downstream of LC14a1 (**Fig. S7**). We uncovered a gradient in the input structure for these two cell types, from exclusively binocular to exclusively monocular (**Fig. 4O**). Thus, binocular disparity-tuned LC10a and monocular size-tuned LC10a target distinct downstream pathways indicating disparity cues could drive a distinct set of behaviors.

### Silencing LC14a1 impairs distance estimation during social interactions

Given the established role of LC10a in male courtship (Collie et al., 2026; Cowley et al., 2024; Hindmarsh Sten et al., 2021; Ribeiro et al., 2018), we focused on distance-dependent aspects of this behavior (**Fig. 5A**), including circling (Ning et al., 2022), chasing (Agrawal and Dickinson, 2019; Hindmarsh Sten et al., 2021), and singing (Roemschied et al., 2023). We silenced LC14a1 neurons (**Fig. S8**, see **Methods**), to render downstream circuitry sensitive to only monocular cues, thus impairing discrimination of objects at different depths. We tracked posture (Pereira et al., 2022) and segmented song in pairs of male and female flies (Coen et al., 2014) (see **Methods**).

**Figure 5.**
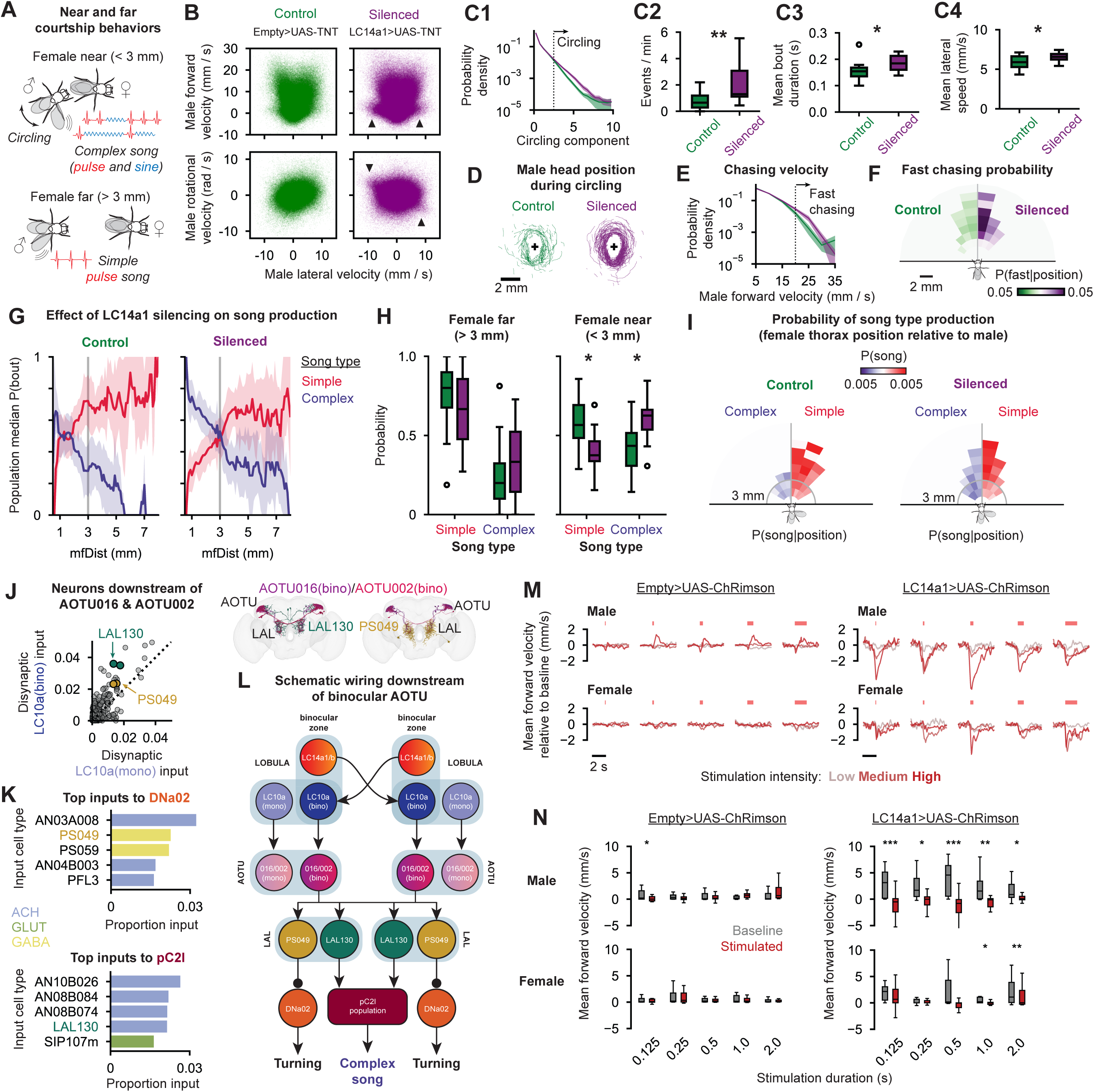
Binocular vision is essential for courtship behavior. A. Male flies pattern their courtship behaviors with distance to females. Males produce complex song and circle around the female when near, and produce simple pulse song when far. B. Distribution of male movement variables in control (left, green) and LC14a1-silenced (right, purple) flies. Top panels: forward velocity vs. lateral velocity; bottom panels: rotational velocity vs. lateral velocity. Black arrowheads indicate increased density of high lateral velocity with opposing rotational velocity. C. Circling behavior is disrupted in LC14a1-silenced flies. C1: projection of male motor variables from B onto the circling principal component where lateral and rotational velocity is anticorrelated (see **Methods** and **Fig. S8**). Line + shading = mean ± s.e.m. probability density across flies. Y axis is log scaled to show differences at extremes of distribution. Circling is defined as times when circling component > 2.5 (dotted line). C2-C4: number of circling bouts (C2), mean circling bout duration (C3) and mean lateral speed during circling (C4) in control (left, green) and LC14a1-silenced (right, purple) flies. * p < 0.05, ** p<0.01, Mann-Whitney U-test. D. Traces of male head position relative to female during circling bouts. Left (green) = control; right (purple) = LC14a1-silenced. Female is oriented facing up with thorax position indicated with+. E. Distribution of male forward velocity during chasing (see **Methods**). Line + shading = mean probability density ± s.e.m. across flies. LC14a1-silenced flies (purple) show more fast chasing (> 20 mm / s, dotted line). Y axis is log scaled to show differences at extremes of distribution. F. Female thorax position during fast chasing. Shading indicates population median probability of male forward velocity > 20 mm / s when female is in different parts of visual field for control (left, green) and LC14a1-silenced (right, purple) flies. G. Song type production relative to distance. Probability of song bout choice with respect to distance for simple pulse song (red) and complex song (blue). Left: control flies; right: LC14a1- sienced flies. Line and shading indicate median and median absolute deviation across files. Gray line indicates near-far context switch (3 mm). H. Silencing LC14a1 increases complex song production. Probability of choosing simple pulse vs complex song when far (left) and near (right) for control and silenced flies. * p < 0.05. I. Probability of simple pulse and complex song production to females in different parts of the visual field. Histogram of female position relative to male for control (left) and LC14a1- silenced (right) flies. Shading indicates population median probability of producing simple pulse (red) or complex (purple) song bouts at a given position. J. Neurons downstream of the binocular AOTU. Left: each point is a neuron downstream of AOTU016 and/or AOTU002, showing total input from monocular and binocular LC10a within two hops. Right: morphology of LAL130 (teal) and PS049 (yellow) and presynaptically connected binocular AOTU016 (purple) and AOTU002 (pink). K. Top inputs to DNa02 (top) and pC2l (bottom). Mean proportion of input to all neurons of the indicated cell type. The pC2l population consists of multiple cell types (see **Methods**). L. Schematic summary of binocular sensorimotor pathways. LC10a neurons viewing the binocular visual field receive input from the contralateral lobula via LC14a1 and LC14b. Binocular LC10a neurons connect to a subset of AOTU016 and AOTU002 neurons, which in turn connect to LAL130 and PS049. LAL130 drives complex song via pC2l and PS049 inhibits DNa02- mediated turning. M. Optogenetic stimulation of LC14a1 decreases forward velocity. Top left: control males (n=20); bottom left: control females (n=20); top right: LC14a1-Chrimson males (n=20); bottom right: LC14a1-Chrimson females (n=18). Each trace shows mean change in forward velocity relative to baseline (2 s period prior to stimulation) for three stimulation intensities and five stimulation durations (15 stimulations per fly per intensity per duration). N. Forward velocity before (grey) and during (red) high intensity optogenetic stimulation over all flies. Left = control; right = LC14a1-Chrimson; top = male; bottom = female. *p < 0.05; **p < 0.01; ***p < 0.001, FDR corrected Wilcoxon signed-rank test.

LC14a1-silenced males show increases in lateral speed, and alterations in the correlation between rotational speed and lateral speed (**Fig. 5B**). These phenotypes suggest changes in circling behavior (known to rely on the coordination of lateral and rotational speeds at a fixed distance from the female (Ning et al., 2022)) (**Fig. S8**). Indeed, we find that circling is increased in LC14a1 silenced males (**Fig. 5C, Supp. Video 1**), and that silenced males circle at more variable distances from the female (**Fig. 5D**). Normally, males initiate circling when located in front of or at the side of the female, and utilize circling to move behind her for subsequent behaviors (Ning et al., 2022).

LC14a1 silenced males often fail to transition out of circling behavior. While overall, LC14a1 silenced males are able to pursue females, we found that they chase females at higher speed (**Fig. 5E**), particularly when she is centered and distant (**Fig. 5F**), indicating that males also use binocular cues to regulate aspects of chasing. We also uncovered defects in song pattering. Control flies switch from simple pulse song (produced at greater distances) to complex song (produced at near distances) at ∼3 mm (from the male head to the female thorax, corresponding to ∼2 mm from the male head to the female abdomen), whereas silenced flies continue to sing complex song at greater distances (**Fig. 5G-I**).

We analyzed the male CNS connectome (Berg et al., 2026) to uncover putative circuit mechanisms for these phenotypes (**Fig. 5J-L**), and focused on neurons downstream of AOTU016 and AOTU002 (**Fig. 4N**). Among the top downstream targets are PS049 and LAL130 (**Fig 5J, Fig. S9**). PS049 is the strongest inhibitory input to DNa02 (**Fig. 5K**, top). Unilateral activation of DNa02 drives changes in rotational speed (turning), while bilateral activation drives forward walking (Rayshubskiy et al., 2024). This suggests that stimuli that drive LC14a1, which we predict to include other flies centered in the visual field and 2 mm away, should inhibit turning by providing equal inhibition to DNa02 in both hemispheres (**Fig. 5L**). The cessation of circling is one such behavior that requires a drop in rotational speed (**Fig. 5B**). Similarly, LAL130 provides the strongest excitatory input to pC2l from the central brain (**Fig. 5K**, bottom). Activation of pC2l neurons (and their postsynaptic descending neuron targets pIP10 and pMP2 (Lillvis et al., 2024)) drives a switch from simple song to complex song production when males are oriented towards and near females (Roemschied et al., 2023). Thus, this connection provides a mechanism by which binocular distance information can drive the switch from simple pulse to complex song when females are close and centered (**Fig. 5L**). Silencing LC14a1 provides this circuit with only monocular input, causing males to produce complex song when the female is far (**Fig. 5G-I**).

Finally, we activated LC14a1 neurons via csChrimson (Klapoetke et al., 2014) (see **Methods**). This manipulation (in solitary flies) should mimic the appearance of an object in the binocular visual field at all elevations simultaneously. We found that this perturbation caused both male and female flies to slow and walk backwards (**Fig. 5M-N, Fig. S10, Supp. Video 2**), with a stronger effect in males, suggesting again that the behavioral circuitry downstream of LC14a1 is sexually dimorphic.

## Discussion

Our discovery of the circuitry for stereopsis in *Drosophila* relied on a complete synaptic resolution connectome spanning the two optic lobes (Berg et al., 2026; Dorkenwald et al., 2024), on linking this connectome to a 3D model of the eyes (Zhao et al., 2025), and on methods to predict neural tuning from wiring. We show how using wiring alone, one can uncover the circuit basis for a conserved computation. In the fly visual system in particular, given its stereotyped anatomy and organized columnar structure, one can begin with the predicted activity of individual photoreceptors, and follow the wires into the brain to understand how excitatory and inhibitory connections come together to enable the detection of particular features in the environment. In the case of binocular circuitry, this involves loops between the two eyes, and we demonstrate the importance of recurrent circuitry for stereovision. We also show how binocular activity is read out by downstream pathways to enable both stereopsis at a fixed distance as well as social behaviors that rely on distance cues.

We find that LC14 neurons have a particular circuit architecture to enable stereopsis. However, this does not imply that other neurons in the *Drosophila* brain do not also perform binocular computations - if so, this must occur in neurons downstream of the optic lobes (in the central brain), as LC14 neurons are the only optic lobe cell type that spans the two hemispheres with receptive fields localized to the anterior visual field (Kim and Kim, 2026).

### Limitations

Stereopsis depends on the size, shape and location of the receptive fields of visual neurons. Here, we infer these parameters from the connectome using the proportion of input synapses to a neuron as a proxy for connection strength. While the exact contribution of each cell type to the tuning of LC14a1 might not be perfectly captured by our receptive field mapping, the proportion of synapses provides a good first approximation of input strength in flies (Liu et al., 2022).

Our model ignores single neuron biophysics (e.g., receptors, ion channel conductances), which may also be important for establishing the tuning of LC14a1 neurons. Here, we focus on the network wiring that enables binocular integration which, combined with our firing rate model, is sufficient to uncover a mechanism for stereopsis in the fly. Our model provides a mechanism for supra-linear addition of binocular inputs in LC14a1, and a full biophysical model of the circuit would be capable of capturing other non-linear dynamics which can enhance disparity tuning and stereopsis.

Our model includes subtractive inhibition; however, we suspect the inhibition from LMa1 to also have a divisive component since it is mediated by chloride conductance. We also do not model inhibition earlier in the circuit, nor microsaccadic sampling of photoreceptors (Kemppainen et al., 2022), which would shape the exact visual features that LC14a1 neurons are selective for. However, given the similarity in input structure to LC10a (see **Fig 4a**), it is reasonable to assume that LC14a1 detects similar visual features, which includes other flies (Cowley et al., 2024; Ribeiro et al., 2018).

### Feedforward vs recurrent circuits for stereopsis

Stereopsis exists in many diverse taxa, including mammals (Ptito et al., 1991; Samonds et al., 2019; Tanabe et al., 2022; Wheatstone, 1838), amphibians (Collett, 1977), birds (Fox et al., 1977), cephalopods (Feord et al., 2020), and insects (Nityananda et al., 2018; Rossel, 1983), indicating that the computation has evolved multiple times independently (Read, 2021). Prevailing computational models for stereopsis were formulated to explain the observed diversity of disparity tuning responses in the primary visual cortex of cats and monkeys (Ohzawa et al., 1990). In these models, monocularly tuned thalamic neurons synapse onto binocular cortical neurons in a feed- forward circuit. However, binocular feedback from cortex to thalamus exists and could potentially aid with stereopsis (Murphy and Sillito, 1989). Recurrent connections also exist in the cortex that enhance disparity tuning (Samonds et al., 2013). A feed-forward model was previously also assumed to be the basis of stereopsis in insects (O’Keeffe et al., 2022), although recurrent connections between the optic lobes have also been postulated in this species (Rosner et al., 2019). Feed-forward models assume that disparity detection results from multiplication in the dendrites of neurons that receive feedforward monocular input. We find in *Drosophila* that binocular information feeds back to earlier processing layers in the contralateral hemisphere corresponding to a specific binocular disparity, resembling a different circuit architecture for stereopsis that was theorized 50 years ago (Grossberg, 1983, 1976). While recordings in cortical neurons have shaped our understanding of stereovision in mammals (Parker, 2007), in the majority of vertebrates, the optic tectum (superior colliculus) is the primary visual processing center in the central brain, and contains extensive interhemispheric connectivity and binocular responses (Berman et al., 1975; Gebhardt et al., 2019; Knudsen, 1982; Tian et al., 2025). Thus, recurrent connectivity may also be an important neural substrate for stereovision in vertebrate models, particularly in sub-cortical structures that are more likely to receive inputs that were filtered by the earlier visual system to match specific ethologically relevant stimuli.

The recurrent LC14a1 circuitry we uncover in flies may have certain advantages. By connecting neurons across hemispheres, flies achieve binocular integration with fewer total neurons, but the circuit can only compute a single binocular disparity, which flies may be able to adjust subtly by changing the vergence movements of the two eyes (Fenk et al., 2022) (see **Fig S4**). In addition, recurrent circuitry may aid with non-linear binocular computations required for stereopsis, provide stability, and allow longer network time constants for integrating binocular information.

While we find that recurrent connectivity is important for computing disparity in LC14a1, LC10a also integrates information from LC14b in a purely feedforward manner. Thus, we find that both recurrent and feedforward circuits for stereopsis are present in the fly brain. The LC14b-LC10a connection is only present at the extreme edge of the binocular visual hemifield. This allows LC10a, as a population, to encode a fixed binocular disparity over a range of azimuths. Such a circuit may be helpful for the fly to determine when a social partner is at a given distance regardless of location within the binocular zone, and thus drive behaviors, such as switching to complex song (see **Fig. 5**), over a range of azimuths, rather than requiring the female to be perfectly centered in the visual field. Alternatively, the LC14b-LC10a circuit could serve as a transition zone for tracking objects smoothly as they enter the binocular visual field. Such tracking of an object between the monocular and binocular visual field could prevent sudden perceptual jumps for the fly, preventing an object from too suddenly being perceived as “close” when it enters the binocular visual field.

### How flies utilize stereopsis

Our efforts to link stereopsis circuitry to behavior focused on male courtship, which involves behaviors produced in response to conspecifics (of a fixed size) at particular distances. We found that the top targets of LC14a1 are visual projection cell types already implicated in sexually dimorphic behaviors, including LC10a, LC43, and LC31 (Berg et al., 2026; Cowley et al., 2024; Deutsch et al., 2025; Schretter et al., 2025). However, we also find that the core connectivity of the LC14a1 circuit is conserved in both males and females (see **Fig. S1**), so it is likely that stereopsis also supports non sexually-dimorphic behaviors. This could include behaviors like collision avoidance (Tanaka and Clark, 2022; Zabala et al., 2012) or landing (van Breugel and Dickinson, 2012), which also rely on estimates of distance in the binocular visual field.

LC10a has been studied for its role in the tracking and pursuit of females by male flies (Collie et al., 2026; Hindmarsh Sten et al., 2021; Ribeiro et al., 2018). This behavior involves changes in angular velocity to continually reorient towards the female during chasing bouts. Our results indicate that this tracking behavior is driven by the monocular LC10a population (see **Fig. S9**). Prior work presented objects of a fixed angular size on a screen 10’s of mm away from the fly, which we predict to be a poor stimulus for the binocular LC10a population. Silencing LC10a during courtship has the strongest effect on complex song production (Cowley et al., 2024), a behavior known to vary with distance (Roemschied et al., 2023) (see **Fig 5**). Our finding that monocular and binocular LC10a target distinct downstream pathways provides a mechanism by which different populations of the same cell type can control the distinct sensorimotor transformations of tracking and song production, respectively.

If flies dynamically alter the convergence of eyes, then LC14a1 activity would vary as a function of both distance and eye convergence (see **Fig. S4**). Currently, we do not know how eye position information is integrated into the stereopsis circuit, and whether this occurs in the optic lobe or downstream in the central brain (e.g. via corollary discharge, (Falt et al., 2026)). One possibility is that vergence movements of the eyes could be detected in visual circuits via reafference, possibly by LC14a2 neurons (see **Fig. S2**). LC14a2 are only weakly recurrently connected (about half as strong as the recurrent connections between LC14a1) and receive input from local motion detection neurons tuned to progressive motion only (see **Fig. S2**); furthermore, LC14a2 neurons do not have an inhibitory surround to their receptive fields. Thus, this cell type could be driven by the correlated motion in the left and right eyes during vergence movements.

### Predicting function from structure

Predicting the function of neurons based on connectivity alone still remains a major challenge, due to extensive interconnectivity and recurrent loops (Lin et al., 2024) and the fact that first-order monosynaptic inputs to a neuron are not always sufficient to explain function (Bargmann and Marder, 2013; Currier and Clandinin, 2025). We believe that connectomes provide two distinct utilities in neuroscience: first, the connectome is useful for testing hypotheses about how a known computation is realized in a given circuit. This approach is predicated on the finding that combining known connectivity with some neural activity information is sufficient to constrain other free parameters, such as neuronal time constants and synaptic weights, to model the function of a circuit (Beiran and Litwin-Kumar, 2025). This approach is probably best highlighted by work on the *Drosophila* central complex (Hulse et al., 2021; Lyu et al., 2022; Mussells Pires et al., 2024; Westeinde et al., 2024). A second utility is that the connectome is a tool for uncovering the latent computational abilities of a nervous system. In this case, rather than the connectome being used *post hoc* to explain the observed activity within a circuit, it serves *a priori* to make quantitative predictions about the, as of yet not measured, function of a circuit. The latter is important for using a connectome to drive new discoveries. In contrast with emerging connectome-constrained simulation approaches that try to model as many cell types and connections in a circuit as possible (Lappalainen et al., 2024; Özdil et al., 2026; Pugliese et al., 2026), our approach is to abstract away details about specific cell types and focus instead on the latent architecture of the network; in this case, the polysynaptic recurrent connectivity between the binocular zones of the optic lobes, which is relayed by LC14 via multiple different cell types, many of which have not yet been characterized experimentally. Thus, we were able to develop a model for stereopsis in the fly without needing to account for the physiological parameters of every cell type in the network, and without “fishing” for connectivity motifs in the whole connectome that matched preconceived notions for how the circuit ought to look. The connection to LC10a was serendipitous in that it provided a strong testable behavioral hypothesis for the functional role of stereovision in the fly. We anticipate that similar targeted analysis of small microcircuits connected to neurons with known function will be particularly tractable to this approach.

## Methods

### Connectome datasets and code

The female connectome we used for our analysis is the FlyWire Brain Dataset (v783) (Dorkenwald et al., 2024; Schlegel et al., 2024) reconstruction of a Full Adult Female Brain (FAFB) (Zheng et al., 2018); with synapses from Yu et al. (2025) (Yu et al., 2025).

The male connectome we used was male-cns:v1.0 (Berg et al., 2026), a complete reconstruction of an adult male central nervous system (brain and ventral nerve cord).

Both connectomes are from the vinegar fly, *Drosophila melanogaster*, and come from a cross of wild-type strain CS Tully to w1118.

All code for the paper was custom written in Python. Custom code either queried tables downloaded from codex.flywire.ai (FlyWire FAFB data) or the neurprint.janelia.org server (Male CNS data).

### Cell type nomenclature

When comparing male and female connectomes, we use optic lobe cell type nomenclature from Matsliah et al, 2024 (Matsliah et al., 2024) for both male and female connectomes. Corresponding cell type names in the male connectome can be found in Nern et al., 2025 (Nern et al., 2025).

When central brain cell type names differ between male (Berg et al., 2026) and female (Schlegel et al., 2024) connectomes, we use the nomenclature from Berg et al., (2026). In the AOTU, we consider AOTU016 to consist of Berg et al. types AOTU016a, AOTU016b and AOTU016c, and Schlegel et al. types CB0739, CB0007 and CB0359; and we consider AOTU002 to consist of Berg et al. types AOTU002a, AOTU002b and AOTU002c, and Schlegel et al. types CB2070, CB1080 and CB3127.

pC2l consists of the following cell types: AVLP718m, PVLP204m, AVLP567, PVLP210m, AVLP715m, AVLP736m, AVLP735m, AVLP716m, AVLP713m, AVLP737m, PVLP209m, SIP137m_b, SIP137m_a, AVLP738m, SIP108m, SIP109m.

### Connectome analysis

#### Connection weights

We use normalized synaptic counts to estimate the connection strength between pairs of neurons. Specifically, for two neurons, *i* and *j*, we define the connection strength *i*→*j* (*w_ij_*) as the number of synapses neuron *i* makes with neuron *j* (*n_ij_*) divided by the total number of input synapses onto neuron *j* from all neurons (*N_j_*):

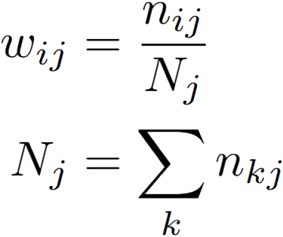

For some analyses, we only consider input to a neuron within specific neuropils, in which case we normalize synaptic counts onto a neuron within those neuropils only, ignoring connections within other neuropils. This is motivated by our understanding that neurons may be compartmentalized or have distinct input and output domains. For example, when assessing the visual inputs to LC9 (a major input to LC14a1 and LC10a, among others), we restricted our analysis to optic lobe neuropils. This is because LC9 receives extensive axo-axonic inputs within its optic glomerulus in the PVLP from central brain neurons. Including synaptic counts from these neurons would skew the visual input statistics to this cell type when compared to optic lobe intrinsic neurons. Given that LC9 dendrites show responses to visual stimuli (Aptekar et al., 2015), we assume visual inputs are sufficient to activate this neuron, while axo-axonic synapses in the central brain may play a role in determining whether that visual signal is propagated to downstream neurons. We acknowledge that, while it is possible that axo-axonic inputs may generate antidromic spikes that propagate back to the optic lobes, such retrograde activity may be less relevant for shaping the visual receptive field of this neuron that its dendritic inputs within the lobula.

Thus, for all analyses in **Figures 1, 2 & 4** (up to investigating the outputs of LC10a in the central brain) we consider only the connectome of the optic lobes (unless otherwise stated), which includes the neuropils of the lamina, medulla, accessory medulla, lobula and lobula plate.

To compute the connectivity between cell types, we summed the input weights from all neurons of a given cell type, *c*, to a focal neuron, *j*, to compute a cell type input weight, *v*_cj_:

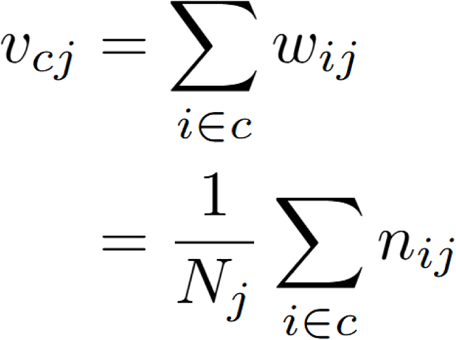

We sorted input types by the mean input they provide to neurons of the focal cell type to obtain the plots in **Figure 1E, 1G, 4C, S1, S2 & S7**. The mean input from one cell type, *c*, to another, *b*, is:

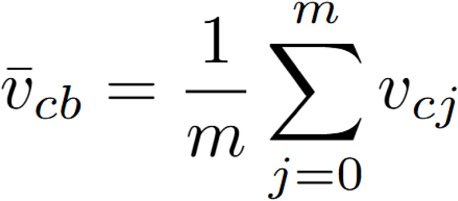

For some figures (e.g. **Figure 1E**), we also show how many neurons of a given type provide input to a target neuron, which helps with interpreting the weight from a given cell type. E.g., the same proportion input to a neuron could come from lots of synapses from a few neurons of a given type (e.g. the LC9 → LC14a1 connection), or fewer per-neuron synapses summed over many neurons of a given type (e.g. the T2 → LC14a1 connection).

#### Excitation and inhibition

Throughout our analysis, we consider excitatory and inhibitory inputs to neurons separately. The effect one neuron has on another depends both on the number of synapses and biophysical properties of those synapses (and neurons). Moreover, the same neurotransmitter can have different effects in different downstream neurons. Comparing the number of synapses from neurons that release the same neurotransmitter is a reasonable first approximation of relative input strength since the underlying channels and molecular pathways that drive changes in membrane potential are likely shared. However, comparing synapse counts between, e.g., cholinergic and GABAergic inputs is less meaningful since the biophysical effects of these neurotransmitters are fundamentally different. In our analysis, we consider all cholinergic neurons to be excitatory and all GABAergic neurons to be inhibitory. Glutamate can be either excitatory or inhibitory in the *Drosophila* brain, but it is usually inhibitory in the optic lobes (Mauss et al., 2015; Molina-Obando et al., 2019) (with the exception of the lamina, whose connections we do not consider in the current study); thus, for simplicity we consider all glutamatergic neurons to be inhibitory. Since the majority of cell types in the optic lobes are either cholinergic or GABAergic, this assumption does not have any major impact on the interpretation of our results, with the exception of the effect of LMa1. LMa1 is predicted to be glutamatergic and thus we believe it to be the primary inhibitory input to LC14a1. This is corroborated by the fact that LC14a1 expresses the glutamate-gated chloride channel, GluClɑ (*Y. Kurmangaliev, personal communication*).

#### Constructing wiring diagrams

To obtain the core wiring diagrams depicted in **Figures 1B** and **4D**, we applied the above formulae to all neurons and cell types of the optic lobes (as defined in (Matsliah et al., 2024)). Wiring in the optic lobes tends to be long-tailed, with a few cell types providing strong input to a given cell type and many cell types providing weak input. In our wiring diagrams, we show all the strongest excitatory feedforward connections (based on proportion of synaptic input) starting with columnar cell types in the medulla. We always show the *n* strongest connections, with *n* chosen for clarity to highlight shared inputs/outputs between cell types, or where there is a notable “jump” in the proportion of input provided by more weakly connected cell types (e.g. T2a and Y3 each provide more than twice as much input to LC14b as the next strongest cell type). Normalizing inhibition is widespread in the optic lobes (Seung, 2024b), so for clarity we also omitted inhibitory cell types that provide normalizing inhibition to medulla and lobula neurons upstream of LC14a1 (specifically, Li18 which provides normalizing inhibition to Tm5f, and LT56 which provides normalizing inhibition to LC9). In our quantitative analyses we consider all connections, including weaker inputs not shown in wiring diagrams.

In **Figure 4L**, we highlight specific pathways we predict contribute to the observed behavioral phenotypes. These connections are explained in the main text.

#### Polysynaptic coupling strength between neurons

When determining polysynaptic coupling strength between neurons (e.g. in **Figure 1K**), we computed what proportion of input can be attributed to an upstream neuron within a given number of synaptic hops. Because of the way we compute connection weights (see above), all the weights onto a given neuron (within a defined set of neuropils) will sum to one. Thus, as we propagated weights back through the connectome, the total propagated synaptic strength always sums to one. To compute connectivity strength between two neurons within n hops, we propagate weights via all paths containing n hops, terminating paths once they reach the target neuron, and summing the resulting synaptic weights. For computing excitatory influence, we only consider cholinergic paths. For inhibitory influence, we consider only the excitatory paths onto inhibitory neurons immediately presynaptic to the starting neuron.

### Receptive field mapping

#### Receptive field mapping

At each step, we normalized input weights to each neuron such that they summed to one (assuming equal strength for all synapses to a given neuron). We considered up to four synaptic hops for excitatory inputs to accommodate the feedforward synaptic chain between strongly-connected cell types Tm20 → Tm5f → Y3 → LC9 → LC14a1, and five hops for inhibitory inputs, allowing us to capture the additional di-synaptic connection from LC9 to LC14a1 via LMa1. Similar to computing *Polynaptic coupling strength* above, we consider all paths through the connectome within n hops, terminating paths once they reach a columnar neuron.

In some panels we restrict receptive field mapping to specific pathways to understand the influence of a given visual feature on a downstream neuron. Specifically, in **Figure 4O** we consider the visual pathways to AOTU016 and AOTU002 neurons via LC10a only, and in **Figure S9** we extend these pathways downstream to LAL130 and PS049.

#### Modeling receptive fields

We modeled the receptive fields of the neurons as 2D gaussian functions with centroids and covariance matrices estimated from the connectome. First, we computed the centroid and covariance matrix for the excitatory and inhibitory receptive fields for each neuron separately as described previously (Seung, 2024a). Briefly, the location of a visual column is defined by the “hexel” basis vectors (denoted ***p*** and ***q***), which are offset by 120°. We mapped hexel coordinates onto a cartesian basis, (***x***, ***y***), with a lattice spacing of 1. Normalizing the weights from visual columns onto each neuron (normalizing excitatory and and inhibitory weights separately), yields a probability distribution:

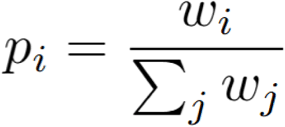

For each neuron, the observed center of this distribution is:

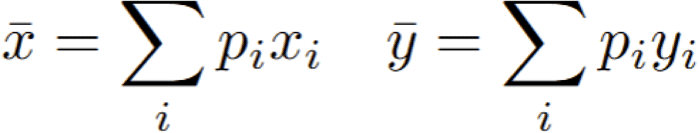

And the observed covariance matrix is:

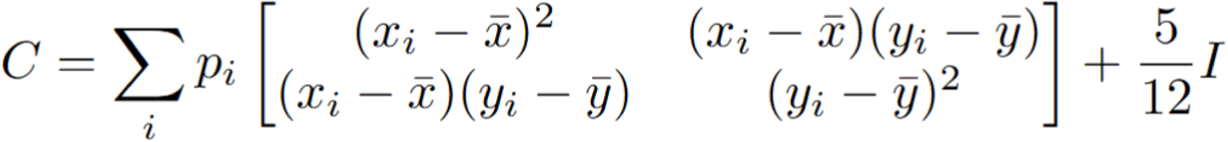

With the 5/12 correction factor accounting for the size of a single hexel.

Since LC14a1 neuron receptive fields lie at the edge of the visual field, the estimate of the centroid and covariance of the input probability distribution is skewed by edge effects (i.e., if the true center of the underlying distribution is at the very edge of the visual field, the observed centroid would be shifted towards the center of the eye). To correct for these edge effects, we modeled the shape of the receptive field to be a truncated gaussian distribution, with the observed half of the distribution defined by a normal vector, *n̂*, pointing towards the center of the eye. This truncated half-gaussian provided a better fit to the underlying connectome weights for LC14a1 than a full gaussian receptive field (**Figure 2I**). Letting *μ* be the observed center (*<u>x</u>*, *<u>y</u>*) and *μ̃* be the corrected center of the distribution:

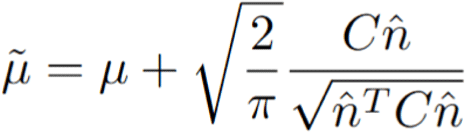

And the corrected covariance, *C̃*, becomes:

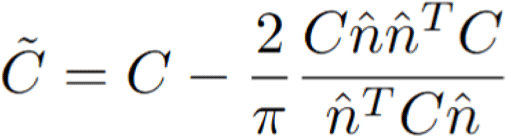

Finally, we modeled the receptive field of each LC14a1 as a 2D gaussian over visual columns with the same generalized variance across neurons (scale factor, *a*), ensuring the receptive fields of each LC14a1 are similar in size. To normalize the size of the receptive fields we scaled the covariances matrices:

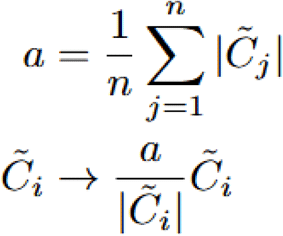

The modeled feedforward input weights from a given visual column, *j*, to a given LC14a1 neuron, *i*, is defined:

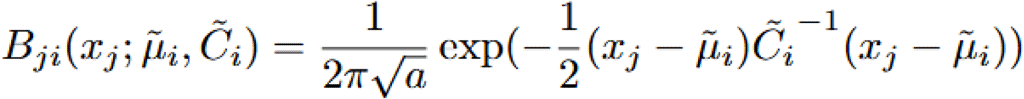

Where *x_j_* denotes the cartesian coordinate of the visual column.

### 3D eye model

We mapped the visual columns from the connectomes to microCT data from Zhao et al. (2025) (Zhao et al., 2025). The microCT data contains the lens and photoreceptor positions for all the ommatidia from an adult female fly. Zhao et al. optimized their fit for the center of the eye, with poorer fit at the edge of the eye. Since these edge points are important for defining the binocular zone, we performed our own fit between the datasets, with the goal of estimating the extent of the contralateral visual hemifield that would be available to a given eye. Performing our own mapping also allowed us to investigate how eye movements would change the size of the binocular zone.

The ommatidia from the microCT data and the visual columns from the connectome each form a hexagonal array that can be described with basis vectors, *p q*, offset by 120°. The number of columns varies across datasets, so we first performed a global fit between the connectome and microCT columns by aligning them along their principal components and rescaling them to have matching variance along each principal component. This performs a 2D global alignment between connectome columns and microCT columns, for which the position and viewing direction of ommatidia are known. We fit a kernel ridge regression model (Python scikit-learn package, 1.7.2) to map the 2D position of microCT columns to a 6D space consisting of the (x, y, z) coordinates of the lenses and (dx, dy, dz) unit vectors of their viewing directions defined by photoreceptor and corresponding lens positions. We then projected the connectome visual columns into the 6D space, renormalizing the viewing direction vectors to have unit length. Ultimately, this provides an estimate for both the physical position of the ommatidium, and its viewing direction, from which a given neural column receives input. To fit the model, we used a fifth degree polynomial kernel, with default values for all other parameters.

Flies can move their photoreceptor array underneath their lenses using two muscles that attach to the retina (Fenk et al., 2022). Contraction and relaxation of these muscles can pull the photoreceptors towards and away from the midline, respectively. Shifting the position of photoreceptors changes the angle at which they intercept light entering through the lenses (which remain fixed). Imaging the location of photoreceptors during behavior shows that flies can perform both conjugate and vergence eye movements (Fenk et al., 2022). Shifting the photoreceptors towards the midline causes them to intercept light from a more lateral angle, effectively causing *divergence*, and shifting them laterally allows them to intercept light from more medial angles, causing *convergence*. The microCT data contain minimal binocular overlap between their ommatidia (Zhao et al., 2025). However, experimental measurements show that movement of the retina can change the ommatidia viewing direction by up to ∼15° (Fenk et al., 2022). Experimental measurements where the flies are free to move their eyes has shown the binocular zone to be up to 30° in azimuth (i.e., with photoreceptors at the innermost edge of the eye seeing up to 15° into the contralateral hemifield) (Kemppainen et al., 2022). We only needed to converge the sightlines of our model ommatidia 5° towards the midline to recover a binocular zone size of 30°, well within the physiological range of movement for the retina.

#### Identifying binocular columns

To identify columns that view the binocular zone, we projected sightlines of ommatidia into space, defining the sightline as the vector from the pseudocone of the rhabdom to the lens. We then computed the shortest distance between sightlines of each pair of photoreceptors taken from the left and right eyes. If this shortest distance spanned a visual angle less than 5° (the interommatidial angle), we considered those ommatidia as intersecting and thus as belonging to the binocular zone of the fly eye. We computed binocular columns for both unconverged and converged states of the eyes (**Figure 2B**). **Figure 2C** shows ommatidia intersections for converged eyes.

### Modeling

We modeled LC14a1 using a rate-based dynamics model, with activity of the *i^th^* neuron denoted *y_i_*. For our model, we let LC14a1 neurons receive input from excitatory and inhibitory presynaptic populations, whose activity is represented by the vectors *x*^+^ and *x***^-^**, respectively. Each LC14a1 neuron has a leakiness determined by the parameter, *α*, excitatory and inhibitory synaptic inputs, and a bias, *β*. We impose a rectified linear unit (ReLU) activation function on the inputs to prevent negative firing rates:

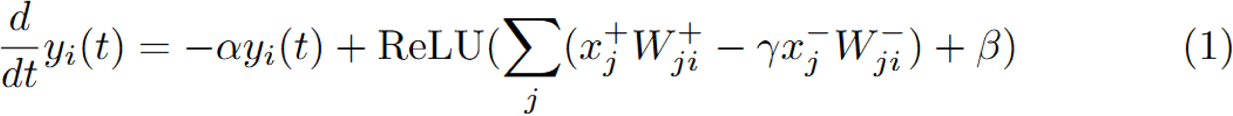

The matrices *W^+^*and *W^-^* contain the input weights from the presynaptic excitatory and inhibitory populations, respectively, and *γ* is a parameter that controls the relative strength of inhibitory inputs.

We let our presynaptic populations be in 1:1 correspondence with neural columns, with their activity modeled as the sum of feedforward input from the retina, *u*, and feedback from LC14a1 whose strength is determined by the weight matrices, *V^+^*and *V^-^*.

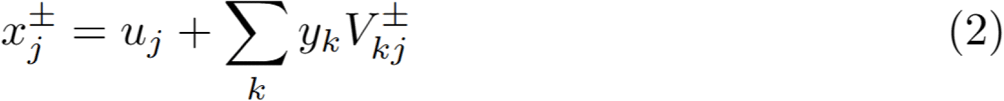

Substituting (2) into (1) yields:

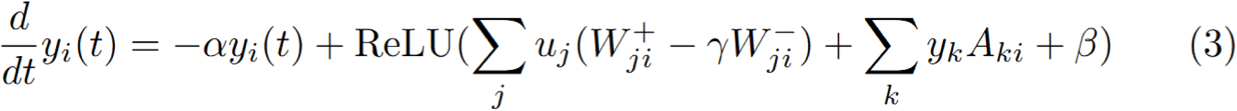

With the elements of *A*:

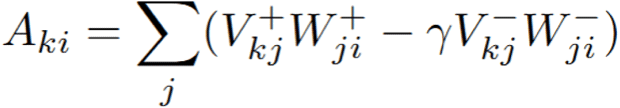

Here, *A* represents the net coupling between pairs of LC14a1 neurons via all paths through the presynaptic population.

We also define a control matrix, *B*, with elements:

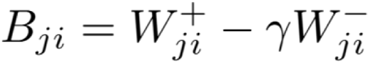

Which maps columnar inputs to each LC14a1 neuron, i.e., the columns of *B* can be thought of as representing the receptive field of each LC14a1 neuron.

Equation (3) is a recurrently connected neural network where all neurons have the same leak and bias, *α* and , with dynamics governed by the weight matrix, *A*, and control matrix, *B*:

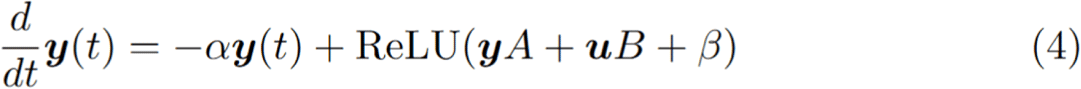

Since the connectome provides a noisy estimate of synaptic connectivity weights, some kind of smoothing or reweighting of synaptic strengths is generally required for firing rate models to produce meaningful dynamics (Duan et al., 2025; Lyu et al., 2022). Since our weight and control matrices, *A* and *B,* have interpretable corollaries in the connectome, we used smoothed estimates for these matrices derived from the connectome in our model.

#### Deriving model weights from the connectome

*A_ki_* is the net coupling between pairs of LC14a1 neurons. We found that LC14a1 neurons are topographically organized and coupled across the midline, with minimal recurrent couplin between LC14a1 neurons on the same side of the brain, and coupling across the midline falling off as a function of distance (see **Figure 1K**). We therefore modeled the connectivity between each pair of LC14a1 neurons across the midline a given distance away as the average connection strength between all pairs of neurons at that given distance. I.e., the modeled connection strength between LC14a1 neurons with the same index on the left and right was equal to the mean connection strength between all neurons with the same index on the left and right, the strength between neurons one index away was equal to the average connection strength between neurons one index away, etc, with the index given in **Figure 1J**. Denoting these averaged connectivity matrices between LC14a1 neurons across the midline, *K*, the weight matrix *A* can be written:

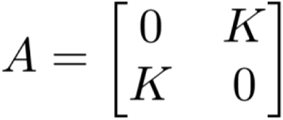

The averaged connectivity matrix, *A*, and its eigendecomposition is shown in **Figure S4**.

With these smoothed weights from the connectome, the final discrete time differential equation governing the dynamics modified from (4) becomes:

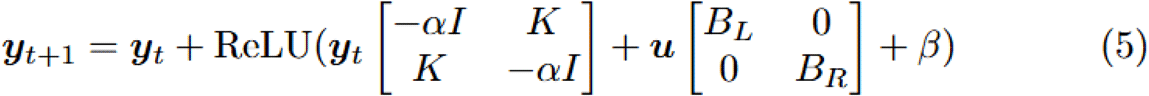

Where *B_L_* and *B_R_* are the receptive fields of the left and right neurons, respectively, modeled as a difference of half-gaussians (see *Modeling receptive fields*, above).

To derive the time traces in **Figure 3D**, we simulated how activity evolved over 300 time steps, with *α* = 0.06, bias (*β*) = −0.015, and initial condition *y*_0_ = 0. For stability, the value of *α* was chosen to be slightly more than the largest eigenvalue of *K*. Our model does not include any scale factors for, e.g., synaptic conductances, so the influence of recurrent feedback within the network is completely determined by the parameter, *α*. The parameter *γ* controls the relative strength of inhibition in the model (see Eq. (3) above). We tested a range of values and for this parameter and ultimately opted for a value of 2.5, reflecting both stronger inhibitory synapses but also accounting for our assumption that activity in neurons providing surround inhibition would likely saturate before a stimulus reaches the full size of their receptive field, whereas neurons providing focal excitation are more likely to be maximally excited by a stimulus that fills their receptive field. In the LC14a1 circuit, we believe inhibition is mediated by chloride conductance, which typically results in inhibitory/excitatory conductance ratios of 7-10 in models of the fly visual system (Ammer et al., 2023), motivating stronger inhibition in our model than would be predicted based on synapse counts alone.

We also wanted to focus on the effects of recurrent excitatory feedback in our model, so we zeroed out recurrent inhibitory feedback (setting the value of *V*^−^ in equation (2) to zero). Since we believe inhibition in the network is mediated by chloride conductance, which is predicted to provide a mix of divisive (shunting) and subtractive inhibition (Chance et al., 2002), we chose not to model the non-linear effects of divisive feedback inhibition and focus instead on the behavior of a more tractable linear model. Since we ignore shunting effects, recurrent inhibition would linearly increase the leakiness of the network relative to recurrent feedback (i.e., reducing the largest eigenvalue of *K*). Since we select *α* based on the eigenvalues of *K,* omitting the recurrent inhibitory feedback from our model does not significantly alter the dynamics.

#### Presenting stimuli to the model eye

The input to our model is a 2D bump of activity on the retina. To compute the shape of this bump for a given stimulus, we projected the sightlines of the model ommatidia into space and computed the angle at which they hit the surface of a stimulus, where the stimulus is a sphere of a defined radius. Specifically, the “activation” of any given visual column is the cosine of the angle between a normal on the surface of the sphere and the ommatidium sightline. This is equal to 1 when the ommatidium views the stimulus head-on and drops smoothly the more oblique the angle at which the ommatidium views the stimulus (0 corresponds to when the sightline is tangent to the stimulus). Our model abstracts away any features of the stimulus that might be computed earlier in the visual system - such as edges, motion or color - and instead we consider only where the object lands in the receptive field of a given neuron.

For **Figure 3C-F**, we presented stationary spherical stimuli of different radii at different elevations, azimuths and distances. We defined the response of a neuron as its steady state firing rate to the stimulus. To perform “blinding” experiments in **Figure 3F**, we measured the response of neurons when providing input only to the one eye. With the exception of **Figure 3F,3M&3N**, which show raw model firing rates, we normalized activity of each neuron to the maximum firing rate that neuron displayed to any given stimulus set.

For **Figure 3G-N**, we presented stimuli of a constant angular size to the model. To achieve this, we scaled the absolute size (in mm) of the stimulus with distance. The angular size of a given stimulus is a function of both distance and azimuth due to the separation between the eyes. To ensure that binocular disparity is the only cue available to the model (rather than size differences when an object is anywhere other than directly in front of the fly), we slightly corrected the stimulus each eye saw to account for interocular distance. We set the radius of a stimulus, *r*, as the scaled sine of the desired stimulus half-angle, *ω*, where 2*ω* represents the full stimulus size (so for a 30° stimulus size, *ω*=15°):

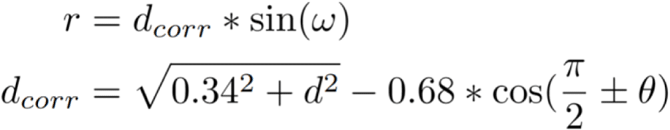

Where 0.68 is the interocular distance in mm, *d* is the distance to the stimulus from the origin (the midpoint between the eyes) and *θ* is the azimuthal position of the stimulus. Here, azimuth is defined in the plane of peak elevation tuning for a given neuron. Object sizes reported in the main text are *2r*.

### Feed-forward model

For the feed-forward (non-recurrent) model in **Figure 3L-N**, we considered an alternative, hypothetical neural architecture where information from the left and right eyes is integrated in a feed-forward network. In this alternative model, downstream neurons integrated activity from the left and right eye according to the receptive fields of pairs of LC14a1 neurons (*B_L_* and *B_R_*):

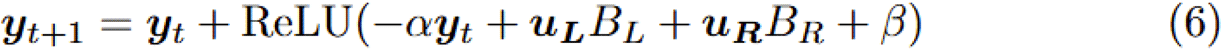

The feed-forward model had the same hyperparameters as the recurrent model (*α* = 0.06, *β* = - 0.015, *γ* = 2.5).

### Behavioral experiments

For all behavioural experiments, we used four- to seven-day-old virgin flies collected from density-controlled bottles seeded with eight males and eight females. Fly bottles were kept at 25 °C and 60% relative humidity. Virgin flies were housed individually and kept in behavioural incubators under a 12 h:12 h light:dark cycling; individual males were paired with a pheromone insensitive and blind (PIBL) female to encourage longer courtship sessions. Experiments were started within 120 min of the incubator lights turning on. Males and females were single and group housed, respectively. Flies were gently loaded into the behavioural chamber before an experiment, using a custom-made aspirator. Chamber lids were painted with Sigmacote (SL2, Sigma-Aldrich) to prevent flies from walking on the ceiling, and kept under a fume hood to dry for at least 50 min before an experiment.

Behavioural chambers were constructed as previously described (Roemschied et al., 2023). Briefly, video was recorded from above the chamber at 60 Hz. Sound was recorded at 10 kHz from 16 particle velocity microphones (Knowles NR-23158-000) tiling the floor of each chamber. Microphones were hand-painted with IR absorbing dye to limit reflection artefacts in recorded videos (Epolin Spectre 160). Temperature was monitored inside each chamber using an analogue thermosensor (Adafruit TMP36).

Song bouts were segmented as described previously (Arthur et al., 2013; Clemens et al., 2018; Roemschied et al., 2023).

#### Silencing experiments

LC14a1 neurons were silenced with genetically-encoded tetanus toxin (TNT) using the split- GAL4/UAS binary expression system. LC14a1 neurons were targeted using the OL0037B split- GAL4 line.

#### Optogenetics experiments

Flies were kept for 4–7 days on regular fly food or food supplemented with all-trans retinal (ATR) at 1 ml ATR solution (100 mM in 95% ethanol) per 100 ml of food. ATR-fed flies were reared in the dark. CsChrimson was activated at 1−205 μW mm−2, using 627-nm LEDs (Luxeon Star).

A fixed stimulus frequency of 1/8 Hz was used for optogenetic neural activation. Stimulus irradiance could take four distinct values (0, 1, 25 and 205 μW mm−2), spanning three orders of magnitude, and stimulus duty cycle could take five distinct values (1/64, 1/32, 1/16, 1/8, and 2/8), and both irradiance and duty cycle were combined in a full factorial design, resulting in 16 distinct blocks that were presented in pseudo-randomized order for 120 s each.

### Behavioral analysis

All analysis was performed using custom written Python code.

#### Tracking

Animal pose was extracted from videos using a deep-learning algorithm, SLEAP (Pereira et al., 2022). We tracked flies with a four-point model consisting of the head, thorax and wings. All SLEAP-tracked data were manually inspected for quality control and any identified tracking errors were corrected.

We defined and computed the following behavioral variables:

- *heading*: unit vector from the thorax to the head
- *mFV* (male forward velocity): velocity (distance in keypoint between adjacent frames) of male thorax in direction of his heading
- *mLV* (male lateral velocity): velocity of the male in the direction perpendicular to his heading
- *mRV* (male rotational velocity): angle between heading vectors vectors in adjacent frames
- *mfDist* (male-female distance): distance from the male’s head to the female’s thorax
- *fAzi* (female azimuth): angle between the male’s heading vector and female’s thorax
- *fFV* (female forward velocity): velocity of the female in direction of the *male’s* heading
- *fFV* (female lateral velocity): velocity of the female in the direction perpendicular to the *male’s* heading
- *mfAlign* (male-female alignment): dot product between the male and female heading vectors
- *fOri* (female orientation): cross product between the male and female heading vectors Derivatives (mFV, mLV, mRV, fFV, fLV) were smoothed with a five frame (83 ms) moving average.

#### Identifying circling bouts

We performed canonical correlation analysis (CCA) between the male’s three movement variables and sensory variables (all data were whitened prior to CCA). The third principal component of the male’s motion corresponded to an anticorrelation between his lateral and rotational velocities. To find circling bouts, we identified all times that the male’s movement projected onto this component was greater than 2.5, the projection onto the first component was greater than −1 (low forward velocity), and the female was within 5 mm and in the front half of the visual field. A circling bout was defined as near consecutive frames (within 100 ms) for which these conditions were true that lasted for more than 100 ms.

#### Identifying chasing

Chasing was defined as times when the male and female’s heading were aligned (within 30°), the male’s forward velocity was greater than 5 mm/s, and the female was in the front half of the visual field and less than 10 mm away. Fast chasing was defined as times the male’s forward velocity was greater than 20 mm/s.

#### Optogenetics

For traces in **Figure 5M**, we extracted six seconds around each stimulation onset for each trial for each fly (two seconds before stimulus onset and four seconds after). For each stimulation, we subtracted the baseline forward velocity, defined as the mean forward velocity in the two seconds preceding stimulation onset. We then smoothed each velocity trace using a 15 frame (250 ms) median filter. The traces shown are the mean velocity over all stimulation events (15 per fly).

For **Figures 5N** and **S10**, for each stimulation event we computed the baseline velocity as above and the mean velocity over the entire stimulation epoch (0.125, 0.25, 0.5, 1 or 2 seconds). We then computed the mean baseline velocity and mean stimulation velocity per fly. We compared mean velocity pre- and during stimulation using a Wilcoxon signed-rank test with false discovery rate correction over all tested conditions (male/female x 3 stimulation intensities x 5 stimulation durations = 30 conditions total).

## Acknowledgements

We thank Sebastian Seung, Ila Fiete, and members of Murthy lab for feedback on the manuscript.

## Author Contributions

DSM and MM designed the study; DSM carried out all data analyses and modeling; YP and IS carried out behavioral data collection; MM obtained funding and supervised the project; DSM and MM wrote the paper.

---

**Figure S1: Supplement to Figure 1.**
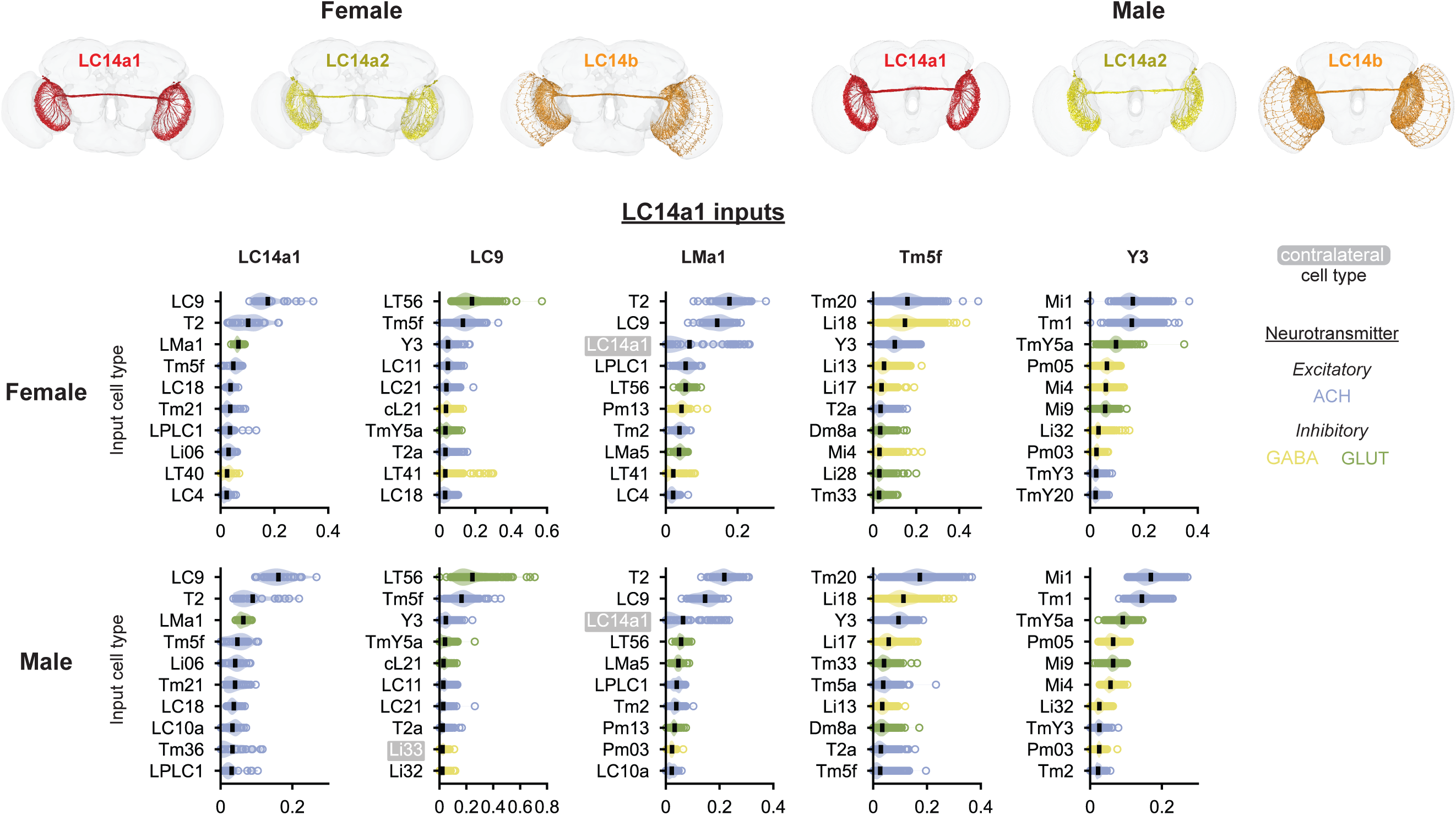
Upper images show all LC14a1, LC14a2 and LC14b neurons in the male and female connectomes. Plots show strongest inputs to LC14a1 and its inputs in female and male connectomes.

**Figure S2: Supplement to Figure 1.**
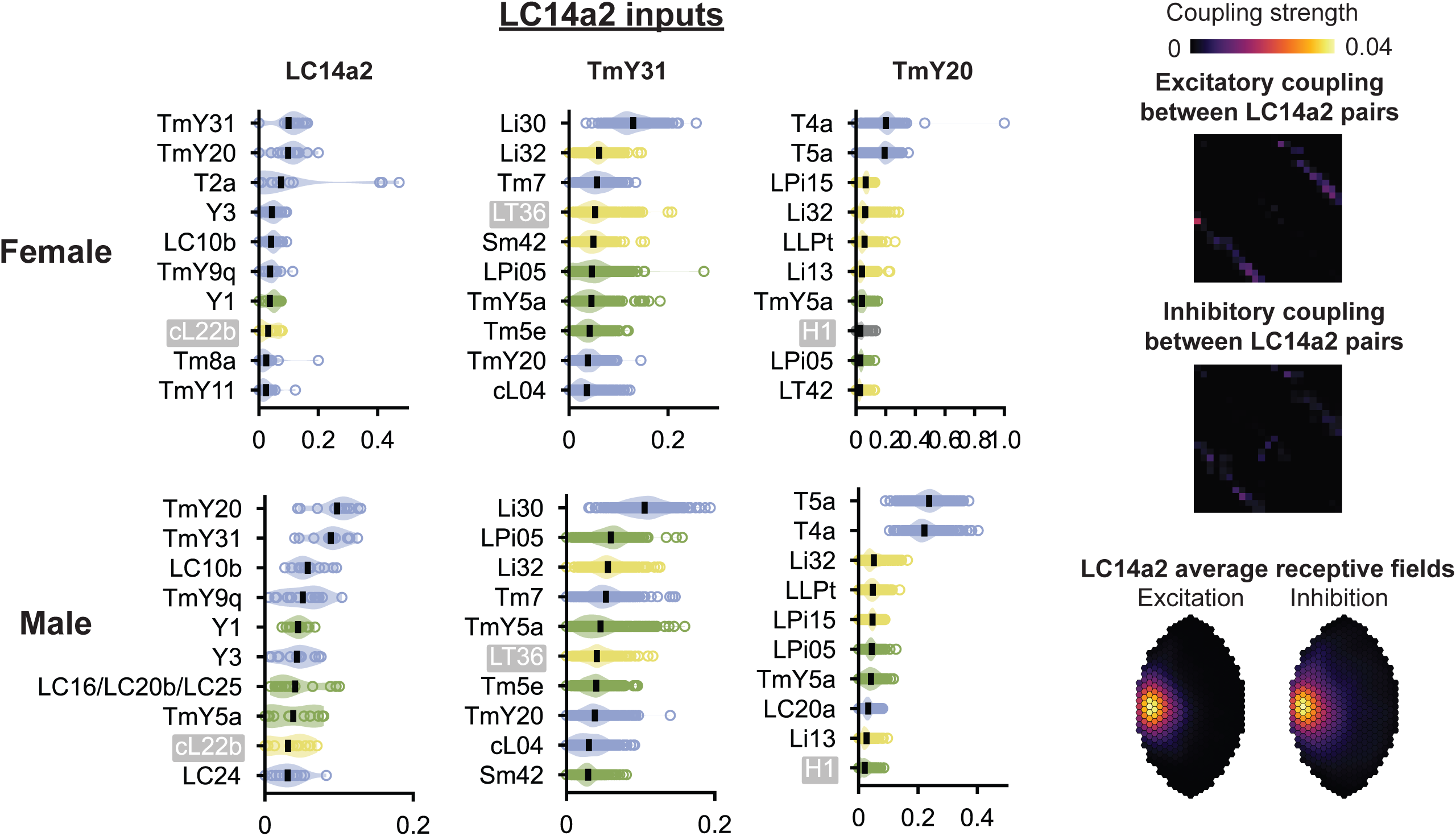
Strongest inputs to LC14a2 and upstream types in male and female connectomes. TmY20 is one of the strongest inputs to LC14a2. T4a and T5a, which respond to progressive (front-to-back) motion, are the two strongest inputs to TmY20. Right panels: LC14a2 is weakly recurrently connected. Lower right panels: average excitatory and inhibitory receptive fields of LC14a2. Excitatory and inhibitory receptive fields overlap, suggesting that LC14a2 does not have a classic center-surround receptive field.

**Figure S3: Supplement to Figure 2.**
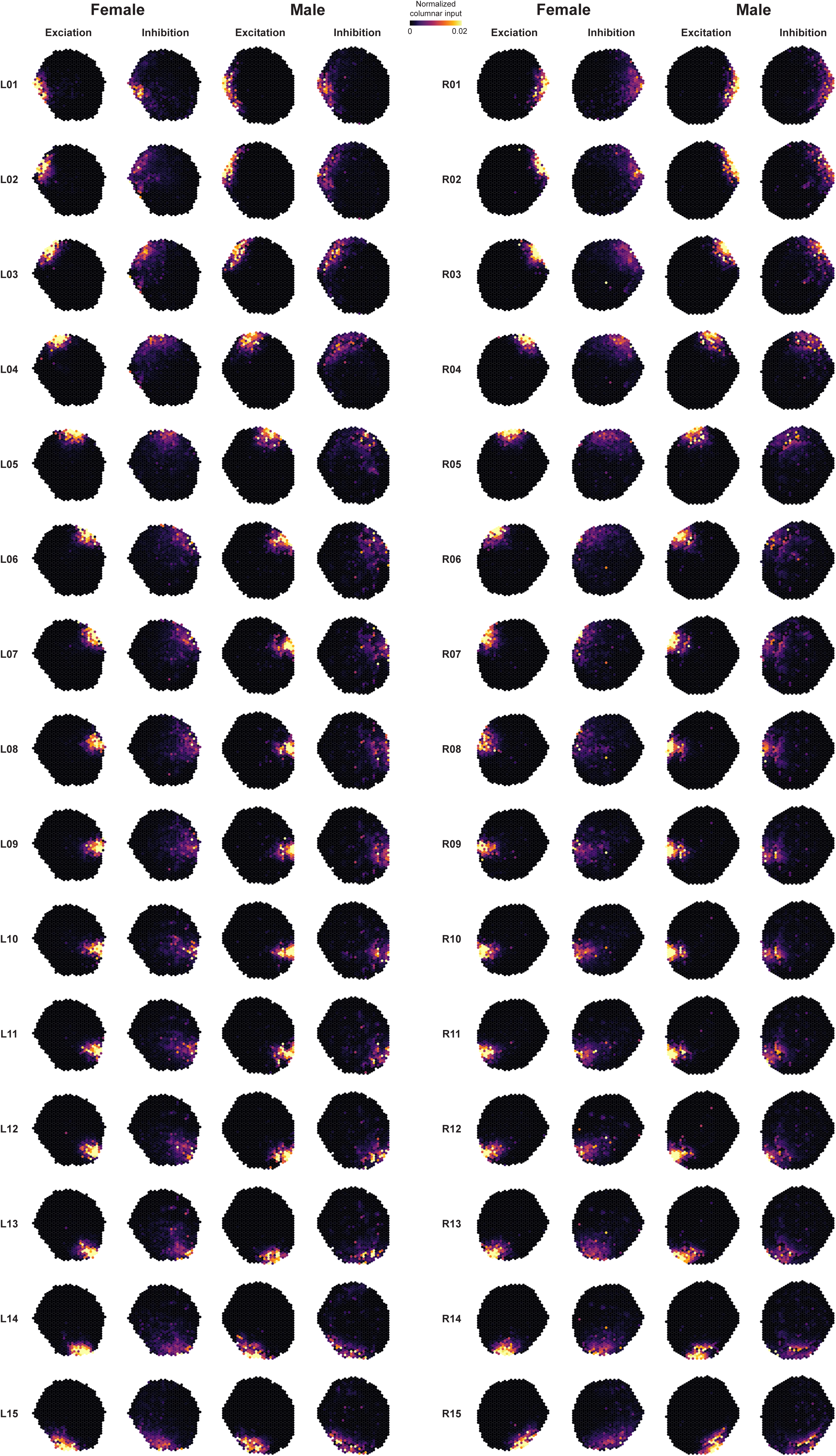
Excitatory and inhibitory receptive fields of all LC14a1 neurons computed from male and female connectomes.

**Figure S4: Supplement to Figure 3.**
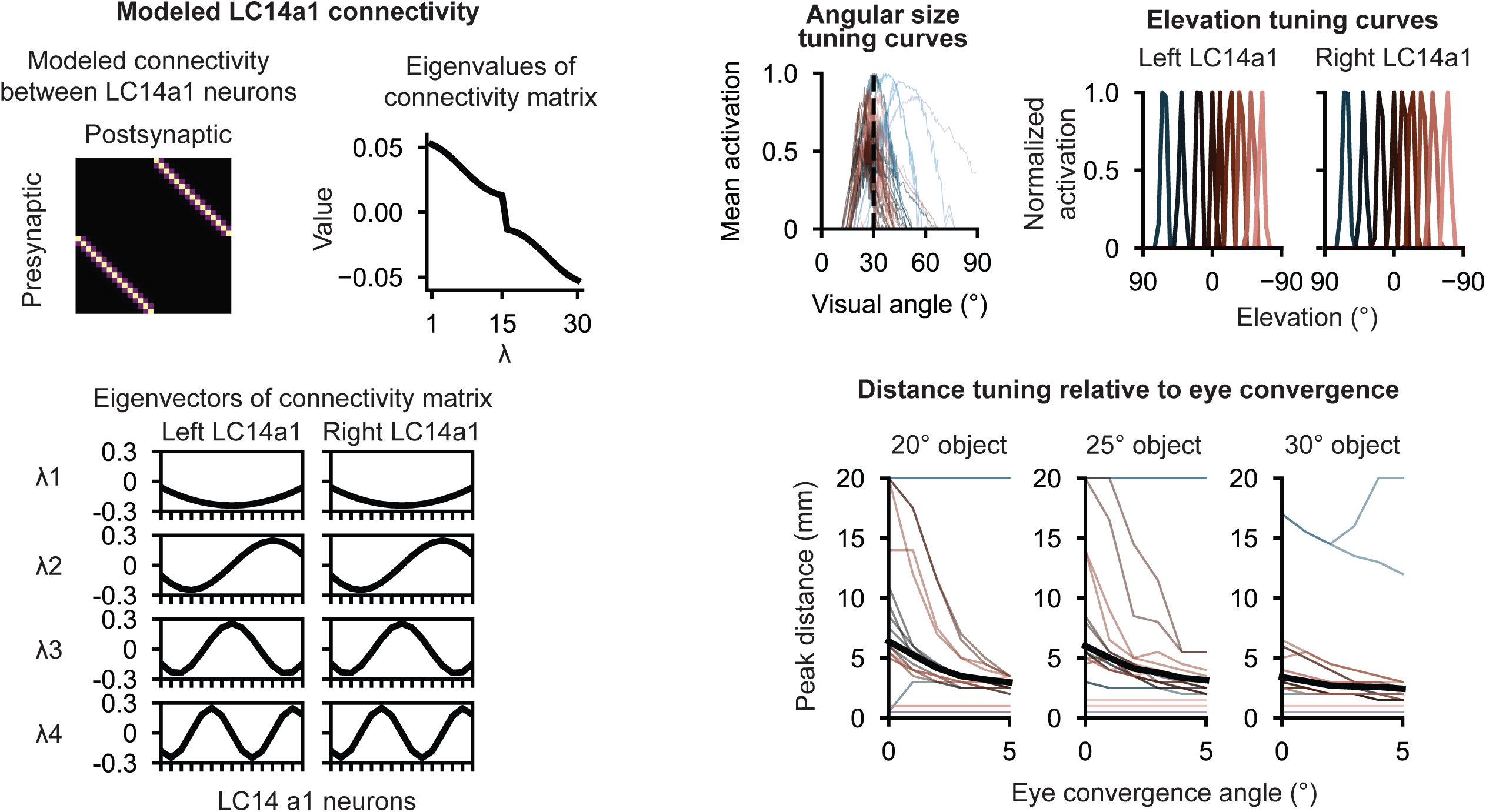
Left: modeled LC14a1 connectivity. Modeled connectivity is obtained by smoothing connections observed in connectome. The eigenvectors of the modeled connectivity matrix form a mirror- symmetric cosine basis across the two optic lobes. Upper right: angular size tuning curves and elevation tuning curves for modeled LC14a1 neurons. Lower right: peak distance tuning to objects of different angular sizes for different eye convergence angles.

**Figure S5: Supplement to Figure 3.**
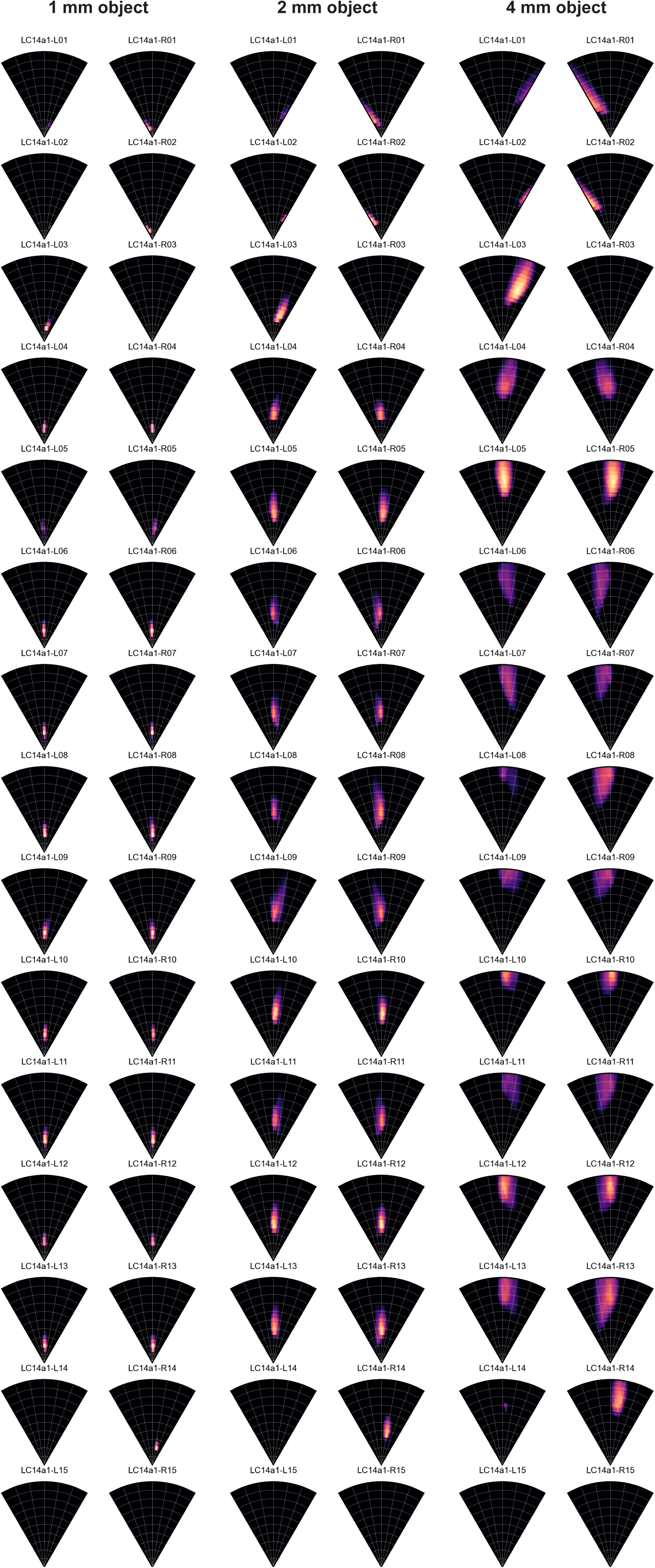
Tuning of all modeled LC14a1 neurons to objects of different sizes (1 mm, 2 mm or 4 mm) at different azimuths and distances, normalized to maximum response across all stimuli tested.

**Figure S6: Supplement to Figure 3.**
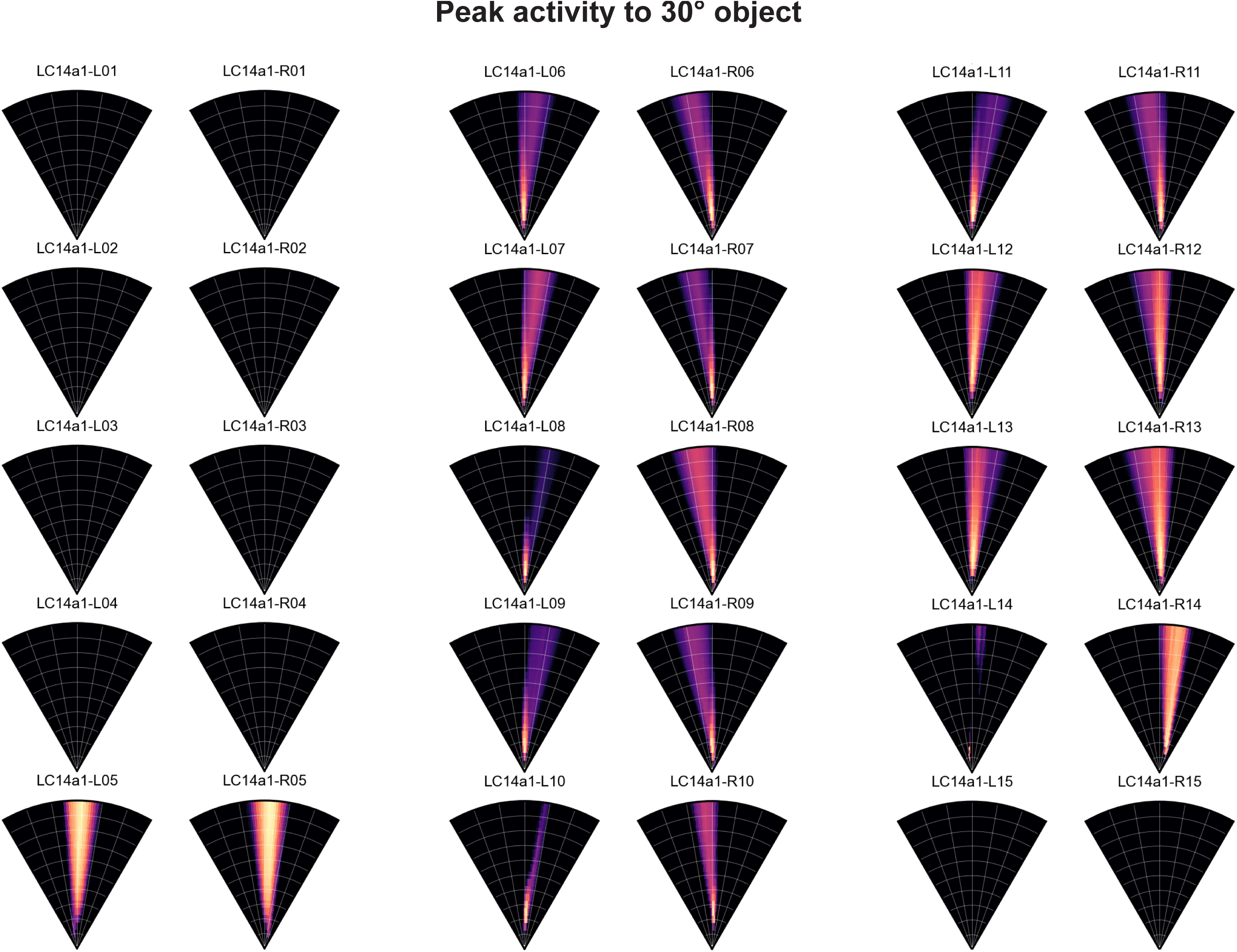
Azimuth and distance tuning of all modeled LC14a1 neurons to a 30° object.

**Figure S7: Supplement to Figure 4.**
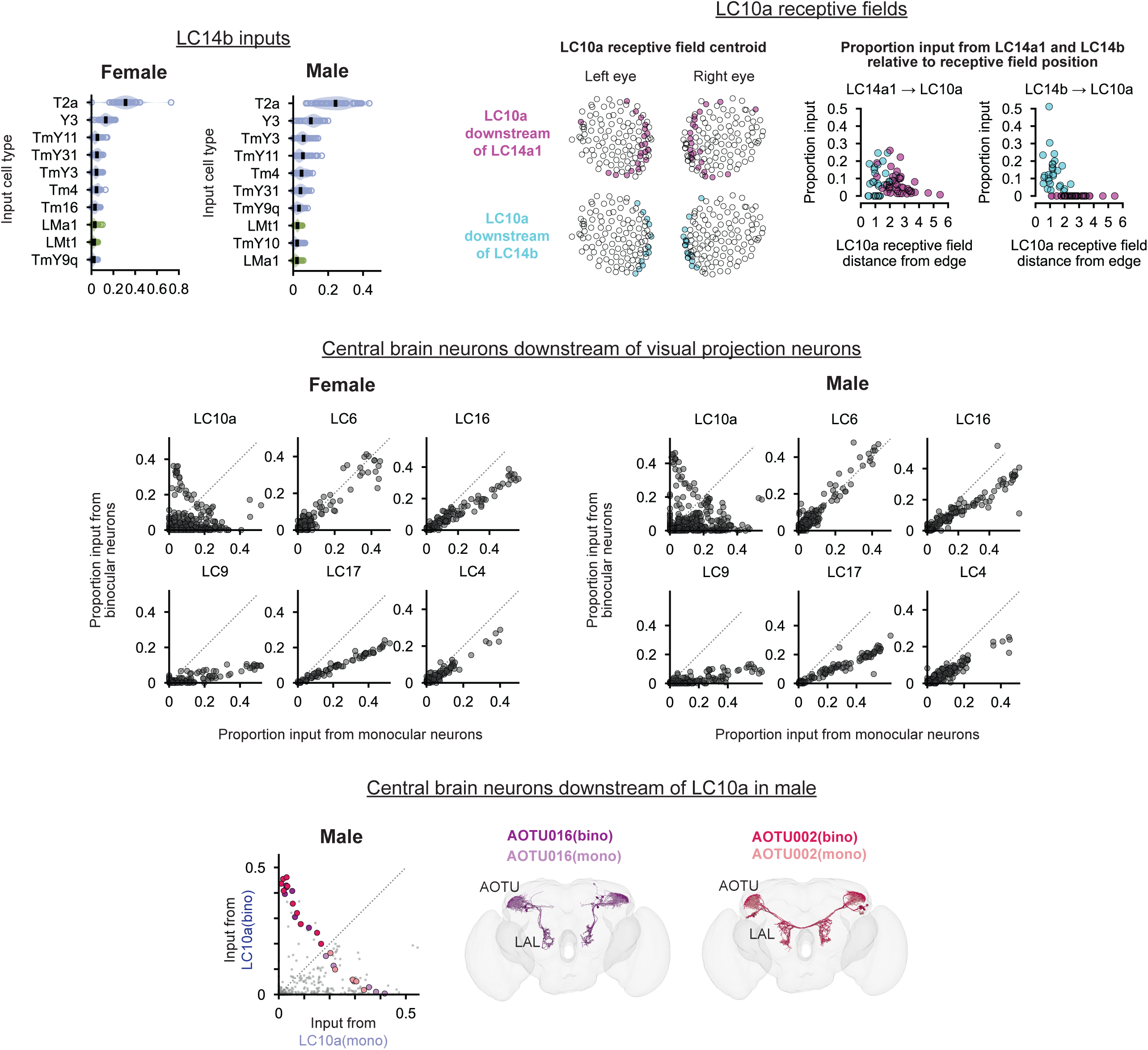
Upper left: strongest inputs to LC14b in males and females. Upper center: centroid of excitatory receptive field of LC10a neurons downstream of LC14a1 only (magenta) or LC14b (cyan). Upper right: proportion of input LC10a neurons receive from LC14a1 and LC14b relative to location of their receptive fields relative to the edge of the visual field (in columns). Center: binocularity of neurons downstream of visual projection neurons (VPNs) strongly connected to LC14a1. Each VPN type is divided into “monocular” and “binocular” populations based on whether it receives input from any subtype of LC14. We then compute what proportion of the total input to a downstream neuron comes from the monocular vs. binocular neurons of the given VPN type. For all VPNs investigated, except LC10a, there is no downstream population that selectively receives strong input preferentially from binocular neurons. Bottom: neurons downstream of monocular and binocular LC10a in the male connectome. AOTU016 (purple) and AOTU002 (pink) neurons are highlighted.

**Figure S8: Supplement to Figure 5.**
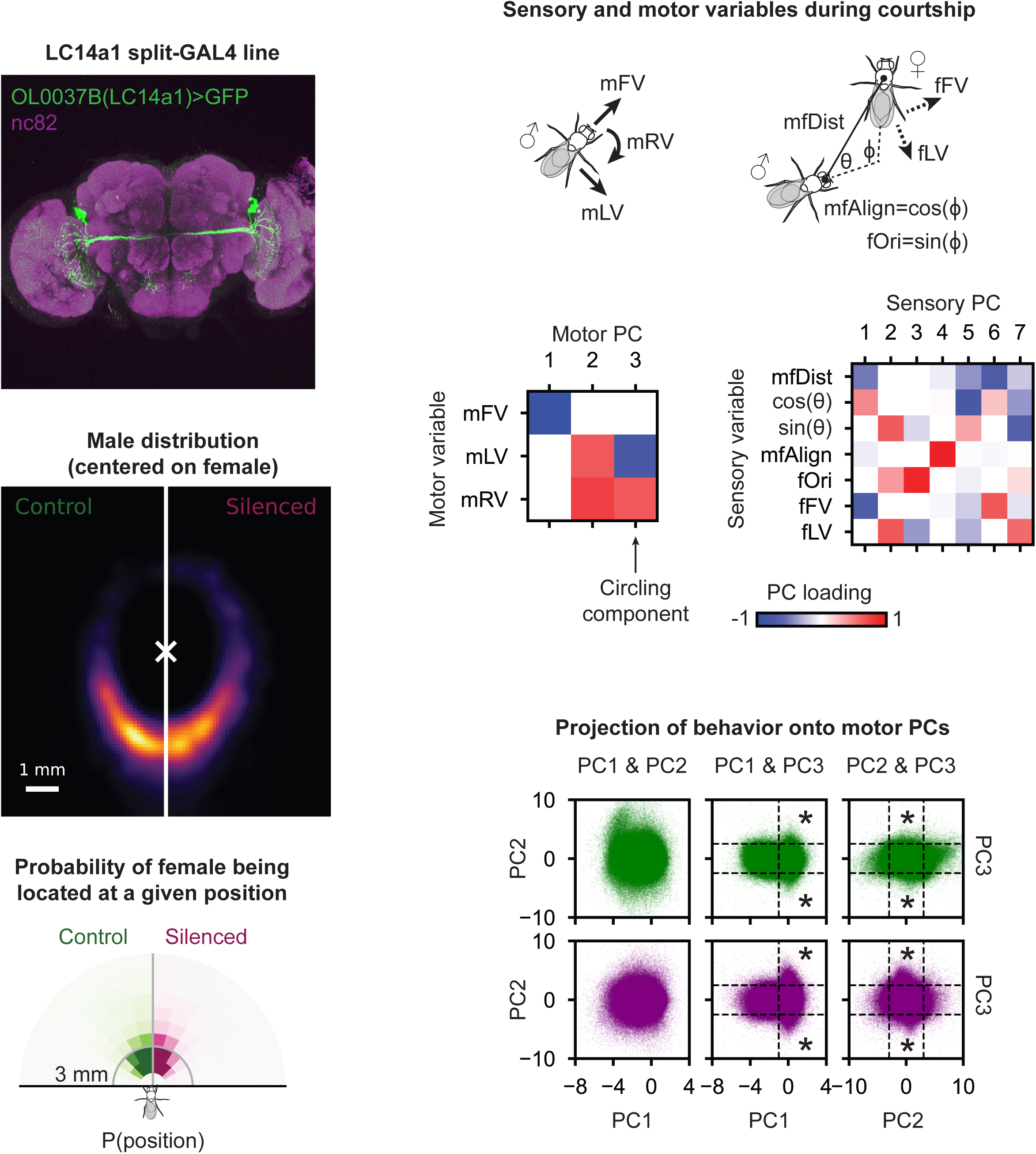
Upper left: confocal image (acquired in Murthy lab) of LC14a1 split-GAL4 line. Lower left: distribution of male around female (centered and facing up, indicated by white x) for control and LC14a1-silenced flies. Lower panel shows histogram of female thorax position relative to male. Upper right: variables extracted from SLEAP tracks in behavioral experiments. mFV: male forward velocity (velocity of male in direction of his heading); mLV: male lateral velocity (velocity of the male in the direction perpendicular to his heading); mRV: male rotational velocity; mfDist: distance from male head to female thorax; fFV: female forward velocity (velocity of the female in direction of the *male’s* heading); fLV: female lateral velocity (velocity of the female in the direction perpendicular to the *male’s* heading); mfAlign: dot product between the vectors defining the male’s and female’s heading; fOri: female orientation relative to the male (cross product between the flies’ heading vectors). Middle right: canonical correlation analysis between the male’s motor variables (mFV, mLV, mRV) and sensory variables (mfDist, azimuthal location (θ), mfAlign, fOri, fFV, fLV), which provides motor and sensory principal components (PCs). Maximum correlation structure is shared between motor PCs and corresponding sensory PCs. Lower right: projection of male’s behavior onto motor PCs. Asterisks indicate region of behavioral space corresponding to circling behavior.

**Figure S9.**
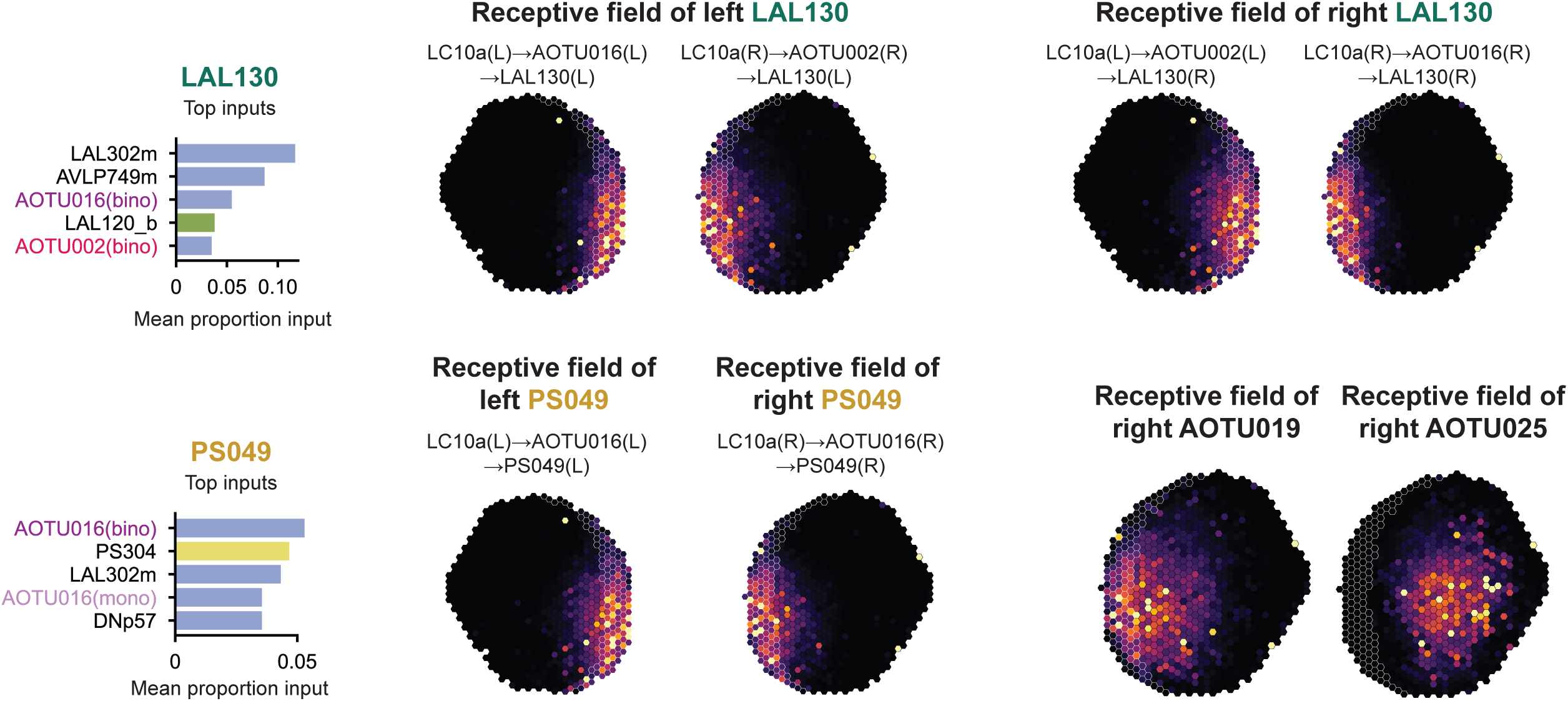
Inputs to LAL130 and PS049. Top: the binocular population of AOTU16 and AOTU002 are among the strongest inputs to LAL130. Maps show receptive fields of the left and right LAL130 computed through the indicated pathway. Bottom left: Binocular AOTU016 are the strongest input to PS049. Maps show receptive fields of PS049 as computed through the indicated pathway. Bottom right: receptive fields of AOTU019 and AOTU025, which drive orienting towards females (Collie et al., 2026).

**Figure S10.**
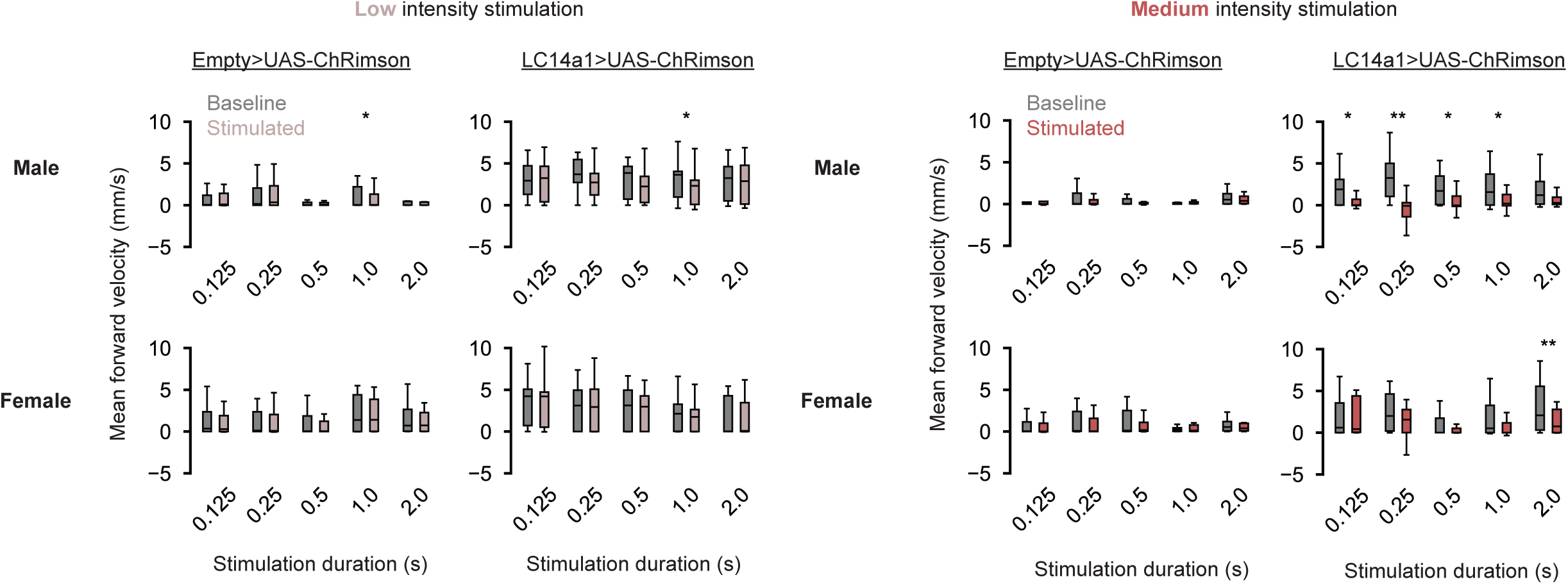
Male and female forward velocity before (grey) and during (red) optogenetic stimulation of LC14a1 at low and medium intensities.

**Supplemental video 1:** example circling behavior in control and LC14a1-silenced flies, showing three examples of each genotype.

**Supplemental video 2:** optogenetic stimulation of LC14a1. Each clip shows one example of a stimulation for each fly in the dataset.

